# Cryo-EM provides insight into how the *Staphylococcus aureus* IsdH receptor removes hemin from the hemoglobin:haptoglobin complex

**DOI:** 10.64898/2026.06.07.730738

**Authors:** Jess Soule, Brendan J. Mahoney, Andrew K. Goring, Joseph A. Loo, Jose A. Rodriguez, Robert T. Clubb

## Abstract

*Staphylococcus aureus* extracts hemin from human hemoglobin (Hb) to overcome host-imposed iron limitation. How it recovers Hb-bound hemin from the hemoglobin:haptoglobin (Hb:Hp) complex, the major circulating form of Hb outside red blood cells, remains unclear. Here we use cryo-electron microscopy, biophysical measurements, and solution kinetics to define how the *S. aureus* IsdH surface receptor extracts hemin from Hb:Hp. A 3.1 Å cryo-EM structure of Hb:Hp bound by full-length IsdH reveals that its N-terminal NEAT domain (N1) anchors it to αHb, whereas its downstream N2N3 extraction unit engages βHb to remove its hemin. The receptor engages Hb:Hp differently than isolated Hb, because N-linked glycans on haptoglobin bias the extraction unit toward βHb, sterically occluding its access to αHb while still permitting engagement by N1. Kinetic assays show that IsdH actively accelerates hemin release from Hb:Hp. Three-dimensional variability analysis indicates that this likely occurs via a dynamic interface in which receptor motions reposition the extraction unit relative to βHb, collectively supporting a model in which IsdH transiently perturbs the F-helix to promote hemin extraction. Alignment of that model with a previously determined CD163:Hb:Hp structure shows how IsdH may disrupt Hb:Hp recognition by macrophage and monocyte CD163 receptors, helping to explain how it may hinder clearance of Hb:Hp from circulation. In aggregate, these results help define the structural basis for hemin extraction from Hb:Hp and how IsdH may subvert receptor-mediated clearance of the Hb:Hp complex.

**Significance:** *Staphylococcus aureus* scavenges hemin from host hemoglobin to proliferate, yet most extracellular hemoglobin is sequestered in hemoglobin:haptoglobin (Hb:Hp) complexes, which are rapidly cleared from circulation. To clarify how bacteria access hemin in this context, we now show that the IsdH surface receptor is structurally adapted to extract hemin from Hb:Hp. Our results indicate that IsdH uses distinct NEAT domains to anchor to αHb and selectively extract hemin from βHb. This βHb selectivity is shaped by haptoglobin N-glycans and enhances microbial access to iron. These findings provide a mechanistic framework for targeting heme acquisition as an anti-virulence strategy.

## Introduction

*Staphylococcus aureus* is a clinically important opportunistic pathogen that causes illnesses ranging from minor skin and soft tissue infections to life-threatening endocarditis, toxic shock syndrome, pneumonia, and septicemia (1, 2). Drug-resistant strains pose a serious public health threat, with methicillin-resistant *S. aureus* (MRSA) causing ∼130,000 deaths globally each year (3–5). To establish infection, *S. aureus* must obtain iron, a critical micronutrient that is limited within the host environment (6, 7). Human hemoglobin (Hb) contains ∼70% of the human body’s iron in the form of heme and is *S. aureus*’s preferred iron source (8, 9). Hb becomes accessible to the bacterium when it is released into the blood plasma upon red blood cell (RBC) lysis, either during natural RBC turnover or in response to bacterial hemolysins (10). To capture its iron, *S. aureus* must overcome host nutritional immunity mechanisms that remove Hb and heme from circulation. Hb is primarily cleared from circulation via the host glycoprotein haptoglobin (Hp), forming Hb:Hp complexes that are removed by CD163-expressing macrophages and monocytes, and, to a lesser extent, through direct CD163-mediated endocytosis (11–15). Free heme is instead cleared by hemopexin and, secondarily, by serum albumin (16, 17). Understanding how *S. aureus* overcomes these clearance systems could lead to new therapeutic strategies to combat lethal infections.

*S. aureus* captures heme from Hb using iron-regulated surface determinant (Isd) proteins (18). Within RBCs, Hb is a tetrameric protein composed of two α and two β globin subunits (α_2_β_2_), each binding a heme prosthetic group. After RBC lysis *S. aureus* encounters methemoglobin (metHb), an oxidized form of Hb in which its heme molecules contain Fe^3+^ (hemin). In the Isd system, the IsdB and IsdH receptors capture metHb on the microbial surface and remove its hemin (19–23). A hemin-binding affinity gradient then drives sequential hemin relay from these receptors to cell wall-embedded hemoproteins IsdA and IsdC, and ultimately to the membrane-embedded ABC transporter complex (IsdEF) (18, 22, 24–26). Hemin is then pumped into the cytoplasm by the transporter, where it is degraded by oxygenases (IsdG and IsdI) to liberate iron (27). Hemin is removed from Hb and trafficked across the cell wall by NEAr iron Transporter (NEAT) domains within IsdA, IsdB, IsdC, and IsdH (28–30). These compact ∼125-residue modules share a characteristic eight-stranded immunoglobulin-like fold and are specialized for either Hb or heme binding (31–37).

The mechanism through which IsdB and IsdH extract hemin from Hb is well understood (6, 9, 10, 38–43); they have closely related primary sequences and remove hemin from metHb using structurally homologous extraction units (IsdB^N1N2^ and IsdH^N2N3^) (22, 23, 34, 44). Each unit consists of an N-terminal Hb-binding NEAT domain, an α-helical linker (L), and a C-terminal hemin-binding NEAT domain. The receptor extraction units accelerate the rate of hemin release from metHb up to ∼1,000-fold (22, 45, 46) by distorting its F-helices, which harbor the proximal heme iron-coordinating histidine residue (47, 48). Both receptors also exhibit interdomain flexibility, which promotes hemin extraction in IsdH (46, 49). Although IsdB and IsdH share related features, IsdB has been proposed to remove hemin via a stepwise mechanism in which α-globin (αHb) binding precedes β-globin (βHb) engagement (50, 51), whereas IsdH^N2N3^ shows no such binding order (52). Hemin extraction by IsdH^N2N3^ also requires covalent linkage of its NEAT domains, in contrast to IsdB, which retains activity when its domains are supplied *in trans* (53, 54). Recent work further indicates that IsdB, but not IsdH, exhibits catch-bond behavior when binding Hb, potentially enhancing its function under blood-flow shear (51, 52, 55). Although their extraction units are conserved, IsdH captures hemin more slowly, possibly due to its additional N-terminal NEAT domain (N1) linked by a ∼100-residue disordered segment, and these differences in Hb engagement and hemin extraction may contribute to their distinct virulence phenotypes in animal infection models (31, 55–57).

How *S. aureus* binds and removes hemin from the Hb:Hp complex remains unclear. In healthy individuals, the Hb:Hp complex is the predominant circulating form of extracellular Hb in blood plasma (58–60). Humans harbor two Hp alleles (*Hp^1^* and *Hp^2^*), which encode proteins containing one or two N-terminal complement control protein (CCP) domains, respectively, and a C-terminal serine protease (SP) domain that binds Hb αβ dimers (12, 55) (**Fig. S1A,B**). Each CCP domain contains a cysteine that forms a disulfide bond between Hp protomers, generating distinct Hb:Hp quaternary structures in Hp1-1, Hp2-1, and Hp2-2 phenotypes. Hp1-1 yields only dimeric Hp, whereas the extra CCP domain in Hp2 creates a second oligomerization site, producing higher-order Hp2-containing assemblies in Hp2-1 and Hp2-2. IsdH is the only pathogen receptor known to remove hemin from Hb:Hp. It also interferes with Hb:Hp binding to macrophage and monocyte CD163 receptors, which impairs its clearance from the circulation and likely prolongs microbial access to this important iron source (61, 62). In contrast, IsdB appears primarily responsible for extracting hemin from free Hb, as it fails to efficiently remove hemin from Hb:Hp and competes poorly for CD163 binding (48, 61). The mechanism by which IsdH carries out its hemin removal and CD163-blocking functions remains unclear, since previous structural studies of IsdH bound to Hb:Hp used truncated receptor constructs that failed to completely remove hemin from Hb and/or truncated and de-glycosylated Hp constructs (61, 62).

Here we define the structural basis of hemin extraction from the Hb:Hp complex by IsdH. A 3.1 Å cryo-EM structure of full-length IsdH bound to Hb:Hp reveals that the receptor uses its N-terminal N1 NEAT domain to anchor to αHb while its downstream N2N3 extraction unit engages βHb to remove hemin. This βHb selectivity is enforced by N-linked glycans on the Hp serine protease domain, which sterically occlude extraction unit access to αHb. Solution kinetics and three-dimensional variability analysis (3DVA) indicate that IsdH actively accelerates hemin release, likely through a dynamic interface in which receptor motions reposition N3 against βHb and enable transient distortion of the F-helix. Structural analysis further shows that IsdH directly occludes the CD163-binding footprint on Hb:Hp, providing a rationale for how it impairs clearance of this iron source.

## Results

### IsdH rapidly removes half of Hb’s hemin molecules in the Hb:Hp complex

To define how IsdH harvests hemin from Hb:Hp, we measured the kinetics and extent of hemin removal from metHb:Hp by the full-length receptor (all subsequent experiments used metHb:Hp complexes unless otherwise indicated). We first asked how Hp binding to metHb alters its affinity for hemin. Consistent with previous reports (65–67), scavenging experiments with apo-myoglobin showed that Hp binding metHb dramatically impairs hemin release (**Fig. S1C-F**). In marked contrast, rapid hemin capture from Hb:Hp is observed upon addition of apo-IsdH^FL^, an IsdH polypeptide containing both the receptor’s extraction unit and its N-terminal Hb-binding N1 NEAT domain (IsdH residues 86-660) (**Fig. 1A**). Apo-IsdH^FL^ was obtained by removing hemin from recombinantly produced IsdH^FL^ and shown by solution nuclear magnetic resonance (NMR) to be properly folded (**Fig S2**). Stopped-flow mixing of 30-fold molar excess apo-IsdH^FL^ with Hb:Hp^mix^, a pooled mixture of Hb:Hp1-1, Hb:Hp2-1, and Hb:Hp2-2 complexes used to broadly assess hemin extraction, produced rapid UV-Vis spectral changes at 405 nm (**Fig. S3A**). These spectral changes report on hemin transfer, as only small spectral perturbations occur in control experiments when apo-IsdH^FL^ is mixed with carbonmonoxy-Hb:Hp (HbCO:Hp), which cannot release its heme to the receptor (**Fig. S3B**). The transfer reaction kinetics are biphasic with *k_f_*_ast_ and *k*_slow_ rate constants of 2.46 × 10^−1^ ± 0.01 × 10^−1^ s⁻¹ and 5.61 × 10^−4^ ± 0.07 × 10^−4^ s⁻¹ (**Table S1**). The fast phase greatly exceeds the rate of spontaneous hemin dissociation from Hb:Hp, indicating that IsdH^FL^ actively removes hemin from Hb:Hp^mix^ (**Fig. S1D**).

**Figure 1:**
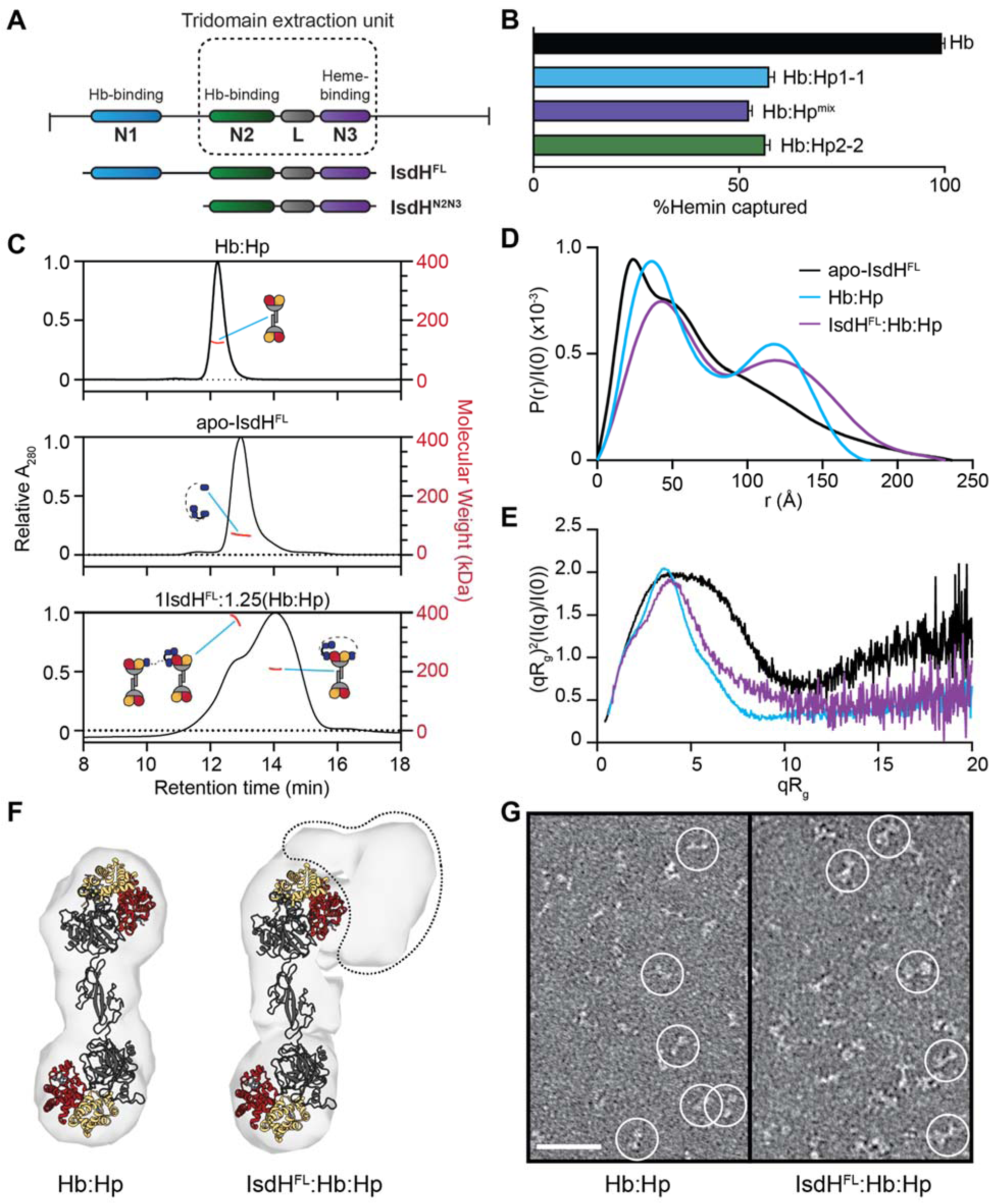
Biochemical and biophysical characterization of the IsdH^FL^:Hb:Hp complex. (**A**) Domain schematic of IsdH and the IsdH constructs used in this study. (**B**) Percent of hemin acquired from Hb (black), Hb:Hp1-1 (blue), Hb:Hp^mix^ (purple), and Hb:Hp2-2 (green). Error bars correspond to values determined by propagation of uncertainty from fits of triplicate measurements. (**C**) SEC-MALS traces of the Hb:Hp1-1 complex (top), apo-IsdH^FL^ (middle), and 1:1.25 apo-IsdH^FL^ and Hb:Hp1-1. (**D**) Distance distribution function P(r) calculated from SEC-SAXS data obtained for apo-IsdH^FL^ (black), Hb:Hp (blue), and the 1:1 IsdH^FL^:(Hb:Hp) complex. (**E**) Dimensionless Kratky plots of apo-IsdH^FL^, Hb:Hp, and the 1:1 IsdH^FL^:(Hb:Hp) complex colored as in (D). (**F**) *Ab-initio* electron density reconstructions of (left) Hb:Hp and (right) IsdH^FL^:Hb:Hp SAXS data using DENSS (63). An existing Hb:Hp model (PDB: 4WJG) (64) is shown docked into each envelope. (Right) Additional density for the IsdH^FL^ receptor is observed at the lobe on the top (circled with a dashed line). (**G**) Negative stain electron micrographs of Hb:Hp (left) alone and (right) in presence of 4-fold excess apo-IsdH^FL^. Scale bar: 50 nm.

Interestingly, IsdH^FL^ only removes ∼50% of the hemin molecules from Hb:Hp^mix^ based on the maximum observed spectral change at 405 nm (^max^ΔA_405_) (**Fig. S3C,D**). Previous studies monitoring hemin removal from metHb showed that the receptor’s extraction unit (IsdH^N2N3^) removes nearly all hemin (23, 44), resulting in ^max^ΔA_405_ of ∼0.3. In contrast, when 30-fold molar excess apo-IsdH^FL^ is mixed with Hb:Hp^mix^ a ^max^ΔA_405_ of only ∼0.15 is reached at the ∼60 min reaction endpoint (**Fig. S3C**). Hemin capture from Hb:Hp^mix^ by IsdH^FL^ plateaus at receptor concentrations below a 30-fold excess, and the UV-Vis spectra of isolated metHb and the Hb:Hp complex are nearly identical (**Fig. S1F**). Thus, the reduced ^max^ΔA_405_ observed for Hb:Hp^mix^ cannot be attributed to incomplete receptor saturation or to Hp-induced changes in the Hb absorbance spectrum. Instead, Hp reduces hemin transfer from metHb to the receptor by ∼50% (**Fig. S3D**). Similar results are obtained for the Hb:Hp1-1 and Hb:Hp2-2 complexes, indicating that fractional hemin transfer is Hp phenotype independent (**Figs. 1B** & **S4**). We conclude that although Hp stabilizes hemin binding to metHb within the Hb:Hp complex, IsdH^FL^ nevertheless actively removes ∼50% of its hemin.

### IsdH undergoes a disordered-to-ordered transition upon engaging Hb:Hp

To characterize the architecture and conformational dynamics of the IsdH^FL^:Hb:Hp complex, we analyzed a sample containing apo-IsdH^FL^ and the Hp1-1 variant pre-complexed to metHb. Hp1-1 was used because it adopts a well-defined dimeric state containing two metHb αβ dimers that is more suitable for structural studies than the heterogeneous higher order oligomers formed by Hp2-1 and Hp2-2 (**Fig. S1A**). The Hb:Hp1-1 complex is used for all subsequent experiments, and is hereafter referred to as Hb:Hp for simplicity. Size-exclusion chromatography of the complex, coupled to multi-angle light scattering detection (SEC-MALS) revealed single peaks with molecular weights (M_w_) of 124 kDa ± 12 kDa and 67.9 kDa ± 0.1 kDa for Hb:Hp and apo-IsdH^FL^, respectively, in close agreement with their expected molecular weights of 150.5 and 66 kDa (**Fig. 1C**). The ternary complex was assembled using sub-stoichiometric apo-IsdH^FL^ relative to Hb:Hp (1:1.25 IsdH^FL^:(Hb:Hp)) to minimize multivalent receptor bridging between Hb:Hp complexes. SEC-MALS of this sample yielded a primary species with a measured molecular weight of 209 ± 2 kDa, consistent with the predicted mass of a 1:1 IsdH^FL^:(Hb:Hp) complex (216 kDa). Its stoichiometry is further supported by native mass spectrometry, which reveals the presence of a 1:1 IsdH^FL^:(Hb:Hp) complex and residual unbound Hb:Hp (**Fig. S5**). A minor species corresponding to two Hb:Hp complexes crosslinked by a single IsdH^FL^ molecule was also detected using SEC-MALS at 381 kDa ± 4 kDa (expected 367 kDa), and is additionally present in the mass spectrum.

To inform on the architecture and conformational dynamics of the ternary complex, we collected small-angle X-ray scattering (SAXS) data for IsdH^FL^:Hb:Hp, and for its Hb:Hp and apo-IsdH^FL^ components. In absence of the receptor, Hb:Hp exhibits a bimodal P(r) function (**Fig. 1D**), consistent with the dumbbell shape observed in crystal structures (64, 68). Overall, the Hb:Hp complex is ordered based on its normalized Kratky plot (**Fig. 1E**), with only modest flexibility indicated by incomplete decay to baseline at high qR_g_ values that presumably originates from motions that reposition its terminal lobes relative to each other (each lobe consists of an αβHb dimer bound to the Hp SP domain). In contrast, isolated apo-IsdH^FL^ is highly flexible, as its normalized Kratky plot remains elevated in the high qR_g_ region. These motions presumably originate from transient repositioning of the N1 domain relative to the N2N3 extraction unit, which are connected by a ∼100 amino acid disordered tether. This is further supported by the receptor’s P(r) function, with substantial probability at large end-to-end distances indicating sampling of extended conformations that maximize displacement of the N1 domain from the extraction unit. Interestingly, binding to Hb:Hp quenches these large-scale dynamics, as evidenced by reduced signal at high qRg in the normalized IsdH^FL^:Hb:Hp Kratky plot relative to that of the isolated receptor. This constraint on receptor motions suggests simultaneous engagement of Hb:Hp by both the receptor’s N1 domain and extraction unit. Furthermore, comparison of the P(r) functions for the IsdH^FL^:Hb:Hp and Hb:Hp complexes reveals that receptor binding only modestly increases D_max_ (from 182 to 232 Å), compatible with full engagement. Stable IsdH^FL^:Hb:Hp complex formation is further supported by *ab-initio* density reconstructions, as both Hb:Hp and IsdH^FL^:Hb:Hp adopt similar dumbbell-shaped structures, with the IsdH^FL^:Hb:Hp complex exhibiting additional density at one end of the complex likely attributable to the receptor (**Fig. 1F**). Consistent with these findings, negative stain electron micrographs reveal similarly shaped Hb:Hp particles that increase in size upon binding by IsdH^FL^ (**Fig. 1G**).

### Cryo-EM structure of the IsdH^FL^:Hb:Hp ternary complex

A cryo-EM structure of the IsdH^FL^:Hb:Hp ternary complex was determined to gain insight into how IsdH^FL^ captures hemin from Hb:Hp. The complex was obtained by incubating 1.25 µM metHb:Hp and 1 µM apo-IsdH^FL^ for ∼30 minutes to allow hemin extraction prior to freezing grids for data collection. 2D projections revealed a globular species consistent with one lobe of an IsdH^FL^:Hb:Hp complex and weak density for the CCP domains and a second lobe that appears blurred because particle alignments were centered on only one end of the complex (**Fig. 2A**). Further processing yielded a 3.1 Å map from a subset of 62,345 particles, and interpretation of that map revealed the IsdH^FL^ NEAT domains bound to a single lobe of Hb:Hp (**Fig. 2B,C**). Signal for the CCP domain-mediated Hp dimer interface and part of the second Hb:Hp lobe are observed in the maps, but are poorly resolved (**Fig. 2B**), reflecting conformational flexibility between the Hb:Hp lobes. This degree of conformational variability is consistent with the modest flexibility identified in the SAXS experiments of the ternary complex and with recent cryo-EM structures of Hb:Hp-containing assemblies (69–73).

**Figure 2:**
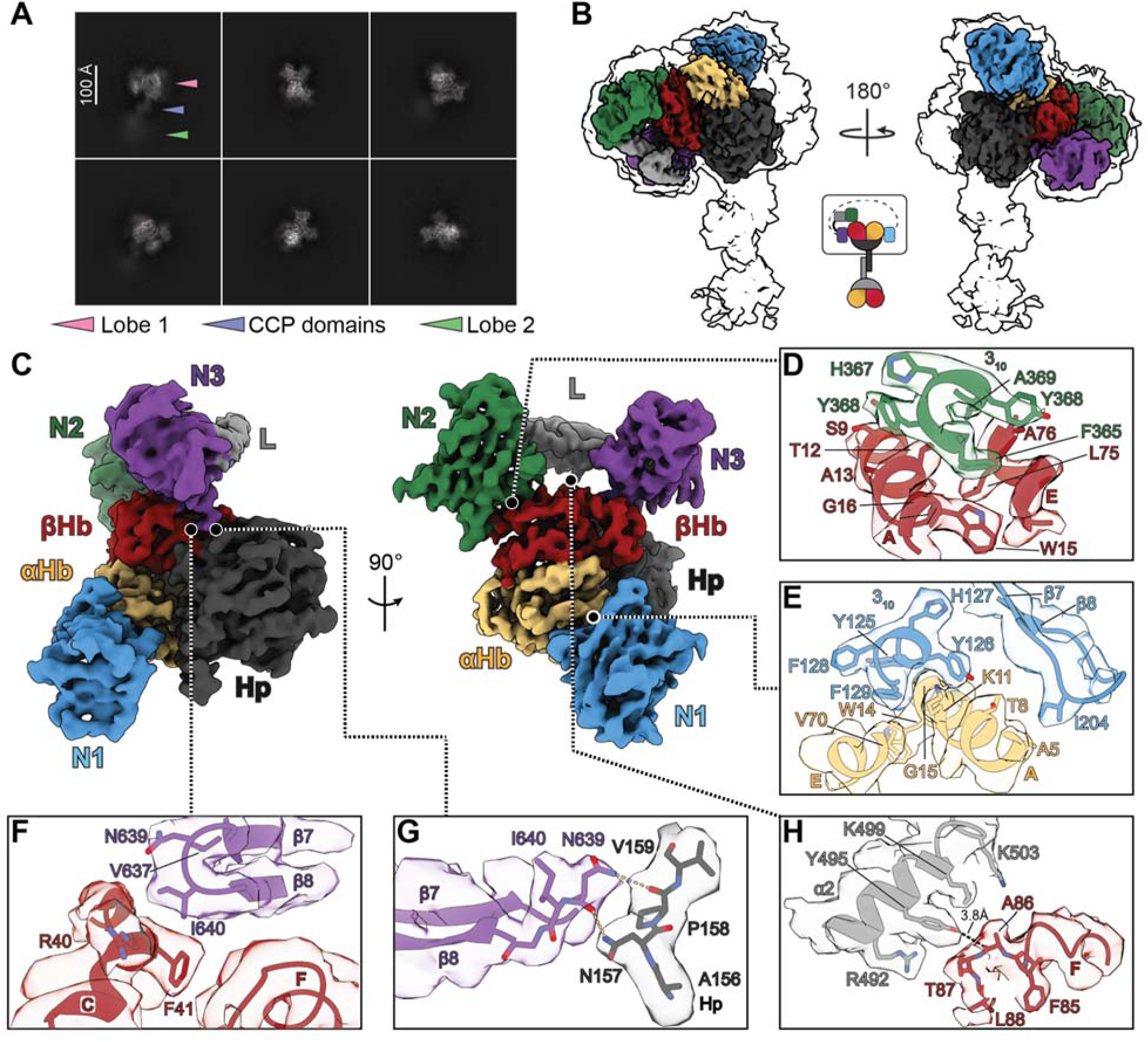
Structure of the IsdH^FL^:Hb:Hp ternary complex. (**A**) 2D classes of the IsdH^FL^:Hb:Hp complex exhibit clear secondary structure features for one lobe (lobe 1: pink arrow). Blurred signal arising from the CCP domains and the second lobe are labelled with blue and green arrows, respectively. (**B**) The IsdH^FL^:Hb:Hp cryo-EM map is represented at high (solid multicolor) and low (outlined transparent) contour. (**C**) Within the 3.1 Å IsdH^FL^:Hb:Hp cryo-EM map, a single Hb:Hp lobe, the IsdH extraction unit (IsdH N2N3), and IsdH N1 are resolved. Interactions between (**D**) IsdH N2 and βHb, (**E**) IsdH N1 and αHb, (**F**) IsdH N3 and βHb, (**G**) IsdH N3 and Hp, and (**H**) IsdH L and βHb. Secondary structure features and amino acid residues are labelled with colors corresponding to their parent protein chain or domain. Hydrogen bonds are depicted as yellow dashed lines, and distances of interest are labelled and shown as a black dashed line.

In the structure, N1 binds to the αHb globin chain, while the extraction unit (N2N3) containing the N2 and N3 NEAT domains and an intervening α-helical linker (L) domain engages βHb and Hp (**Fig. 2C**). The disordered region between N1 and N2 is not resolved. The N1 and N2 domains employ similar structural motifs to anchor IsdH across both Hb globin chains. In N2, a 3_10_-helical lip containing a ^365^FYHYA^369^ motif wedges into a hydrophobic pocket between the βHb A- and E-helices, packing against residues T12, W15, and G16 of the A-helix and L75 and A76 of the E-helix (**Fig. 2D**). N1 mirrors this interaction on αHb, using its own ^125^YYHFF^129^-bearing 3_10_-helix to dock into a similarly positioned hydrophobic pocket defined by W14 and G15 of the A-helix and T67 and V70 of the E-helix (**Fig. 2E**). The N1-αHb interface is reinforced by further contacts to the A-helix: N1 Y126 is oriented to form a cation–π interaction with αHb K11, and I207 in the N1 β7-β8 loop is within van der Waals distance of αHb A5. The IsdH N3 β7-β8 loop, previously implicated in hemin capture (49), contacts both βHb and Hp (**Fig. 2F,G**). Residue I640 forms a hydrophobic interaction with F41 in the βHb C-helix, while N639 participates in hydrogen bonding with Hp, engaging the backbone carbonyl of P158 and the amino group of N157 through its side-chain amide and backbone carbonyl, respectively. Finally, the α-helical L domain of IsdH sits adjacent to the βHb surface, positioning a series of functionally important residues in helix α2 (R492, Y495, K499, and K503) along the F-helix (**Fig. 2H**) (49). Among these, Y495 is proximal to βHb residue A86, with a ∼3.8 Å separation between its hydroxyl group and the A86 backbone carbonyl. In the structure, hemin is bound to αHb (**Fig. 3A,B**), but missing in βHb as a result of transfer to the receptor’s N3 domain (**Fig. 3C**). Notably, the N3-bound hemin is engaged in a hydrogen bond with the βHb distal histidine (H63) suggesting a possible role for this residue in hemin transfer. The hemin-depleted βHb pocket is distorted due to displacement of several adjacent α-helices and F-helix unwinding, which expands the cavity volume from ∼400 Å³ in metHb to ∼512 Å³ in IsdH^FL^:Hb:Hp (**Fig. S6 & S7**).

**Figure 3:**
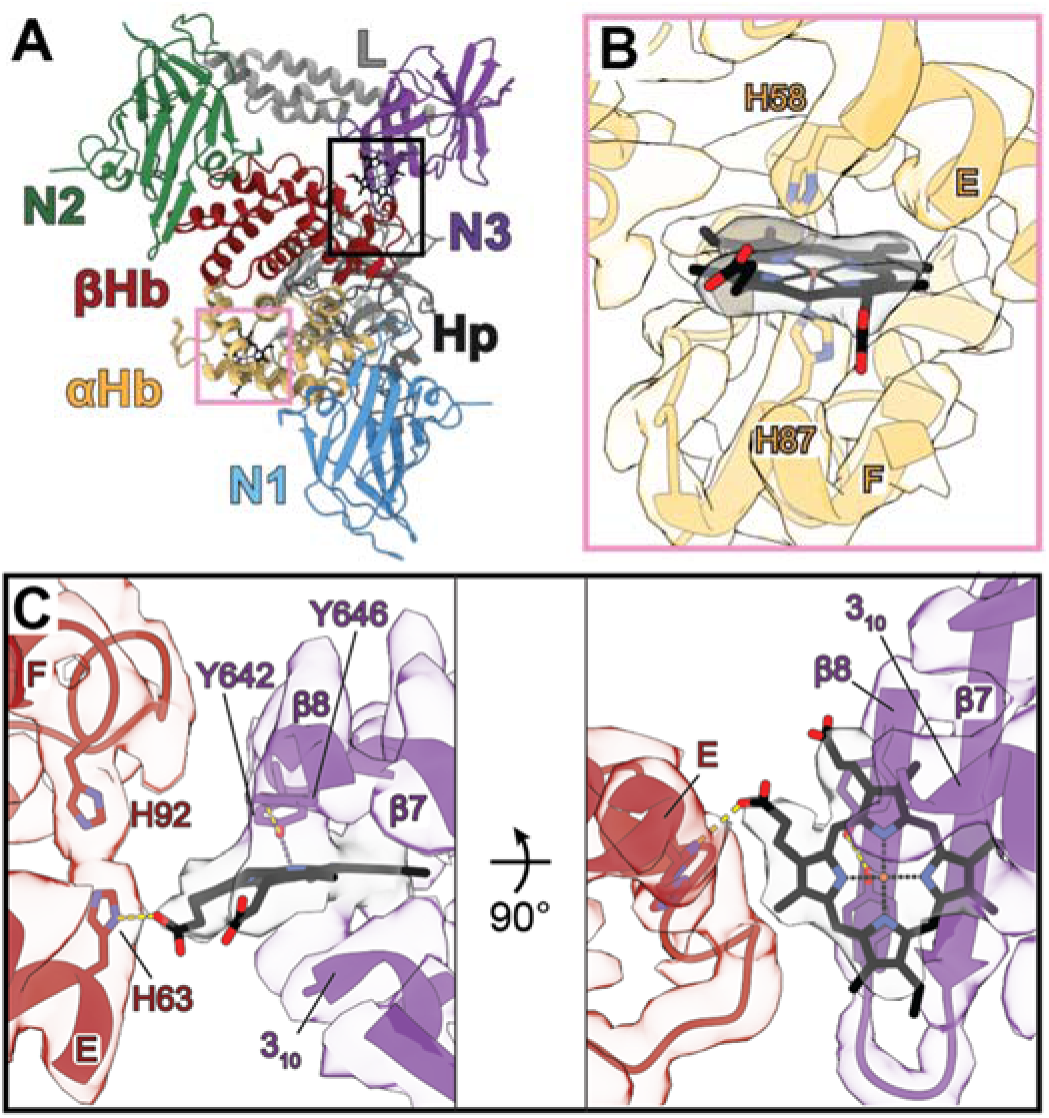
Hemin binding pockets of the IsdH^FL^:Hb:Hp complex. (**A**) Cartoon representation of the IsdH^FL^:Hb:Hp complex colored as in Fig. 2. Hemin molecules are shown as black sticks. The αHb hemin binding pocket is boxed in pink and the βHb and IsdH N3 hemin binding pockets are boxed in black. Hemin is present in the (**B**) αHb heme pocket but absent in (**C**) βHb heme pocket due to transfer to IsdH N3. Secondary structure features and amino acid residues are labelled with colors corresponding to their parent protein chain or domain. Hydrogen bonds are depicted as yellow dashed lines, and coordination bonds are shown as purple dashed lines.

### 3D variability analysis reveals IsdHFL binds Hb:Hp through a flexible interface

Prior NMR and molecular dynamics studies of the IsdH extraction unit bound to Hb suggest that interdomain motions transiently reposition the receptor’s L and N3 domains to engage Hb’s hemin pocket (49, 74). IsdH^FL^:Hb:Hp maps are compatible with these motions, as residues in the IsdH L and N3 domains, and adjacent βHb F-helix exhibit elevated B-factors and lower local resolution (**Fig. S8**). While discrete classification of cryo-EM maps did not reveal distinct conformations, 3DVA in intermediates mode identified a variability component whose particle coordinates approximate a Gaussian distribution (75). Reconstructions of 10 particle subsets spanning this range showed continuous structural heterogeneity in the complex. Maps arising from particle subsets sampling the center of this distribution were suitable for further analysis and accommodated the IsdH^FL^:Hb:Hp model, which was fit into each of the six central maps using Rosetta FastRelax (76). Cα traces of the resulting models were nearly identical overall, except for localized differences in the IsdH L and N3 domains and the βHb F-helix (**Fig. 4A, Mov. S1**). These motions are consistent with the IsdH L domain α2 helix, which contains residues important for hemin extraction (R492 and Y495), engaging and disengaging the βHb F-helix (**Fig. 4B**) (49). These insights, alongside previously reported structures of related intermediates, allow for new mechanistic hypotheses for hemin extraction from IsdH^FL^:Hb:Hp.

**Figure 4:**
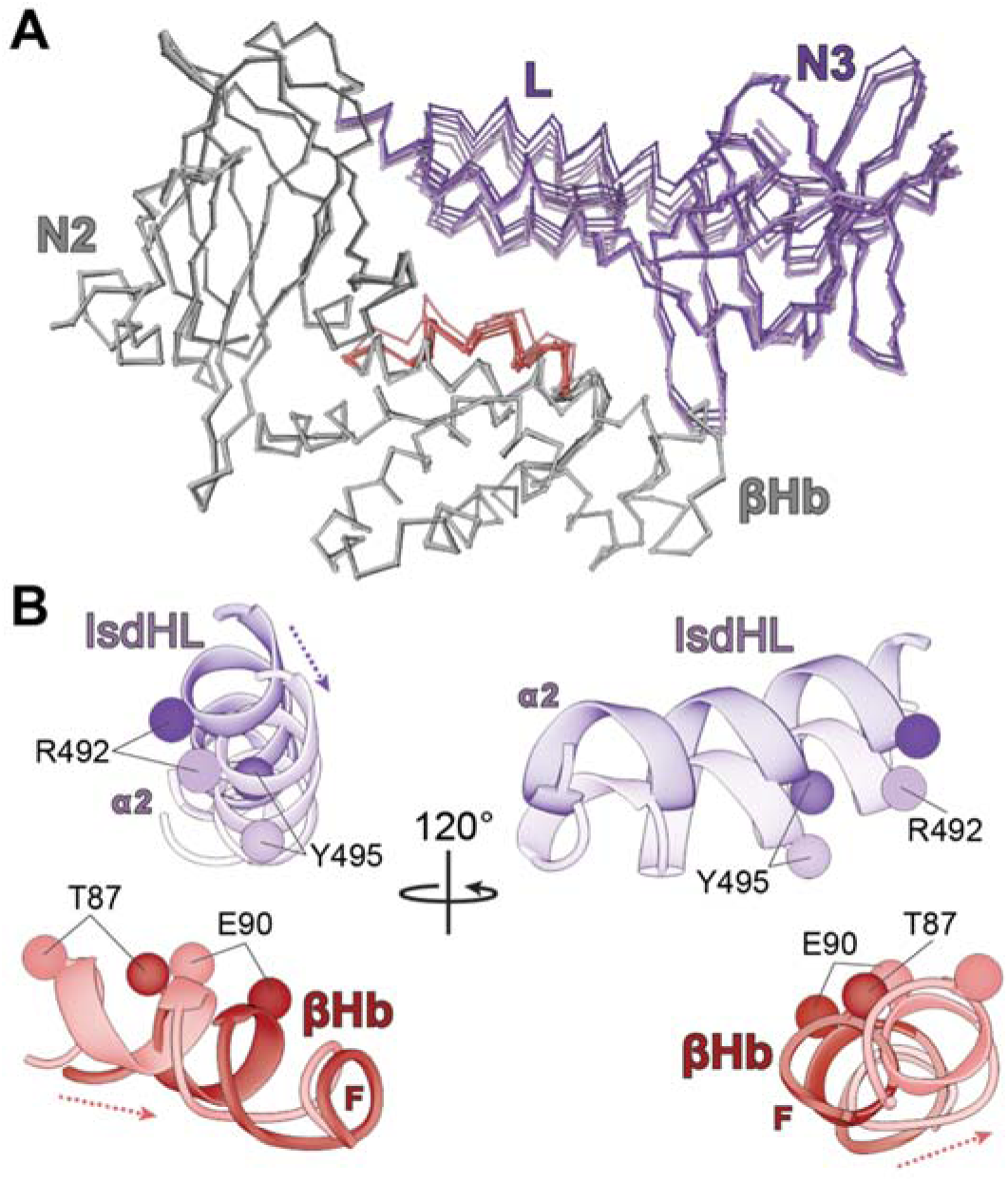
IsdH^FL^ forms a dynamic interface with Hb:Hp. (**A**) Cα traces of six IsdH^FL^:Hb:Hp models generated by Rosetta FastRelax refinement of the consensus structure into maps from 3DVA in intermediates mode. The models are highly similar overall, with conformational differences primarily in the IsdH L and N3 domains (purple shades) and the βHb F-helix (red shades). The IsdH N1 domain, Hp, and αHb are omitted for visual clarity. (**B**) Positional variation between the IsdH L α2 helix and the βHb F-helix at the 3DVA extrema. The dark purple and red models correspond to one extreme while the light purple and red models correspond to the other. Backbone positions of IsdH L residues previously implicated in hemin extraction are shown as purple spheres, and backbone positions of contacting βHb F-helix residues across all available structures are shown as red spheres. Full side chains are not shown due to limited map resolution. Motion trajectories are shown as dashed lines (IsdH: purple, and βHb: red).

### Hp glycosylation targets the extraction unit to βHb

Our combined analyses reveal that IsdH extracts hemin exclusively from βHb within Hb:Hp, in marked contrast to the isolated extraction unit, which removes hemin from both globin subunits in isolated metHb (44). Thus, some unique property of IsdH^FL^ or the Hb:Hp complex must bias hemin extraction toward βHb. Conceivably, binding competition between the N1 domain and extraction unit within IsdH^FL^ could promote globin-specific hemin extraction from βHb. In this scenario, αHb-specific binding by the N1 domain would occlude extraction unit binding, leaving only βHb within the Hb:Hp complex open for hemin removal. To investigate this, we expressed a construct encompassing only the IsdH extraction unit (IsdH^N2N3^; residues 326-660) (**Fig. 1A**) and examined its ability to bind and extract hemin from Hb:Hp. In good agreement with their expected molecular weights, Hb:Hp and IsdH^N2N3^ were measured by SEC-MALS at 124 kDa ± 12 kDa and 38.0 kDa ± 3 kDa, respectively (**Fig. 5A**). Even at 8-fold molar excess, IsdH^N2N3^ predominantly forms a 2:1 IsdH^N2N3^:(Hb:Hp) complex (237 ± 2 kDa; expected 228 kDa), with a minor 4:1 species at 299 ± 3 kDa (expected 305 kDa). These observations and our cryo-EM structure support a model in which IsdH^N2N3^ preferentially engages βHb within the Hb:Hp complex, as equivalent binding to all Hb globin chains (as in isolated metHb) would predominantly yield complexes with 4:1 IsdH^N2N3^:(Hb:Hp) stoichiometry. The notion that IsdH^N2N3^ favors βHb binding within Hb:Hp is further supported by hemin extraction measurements. IsdH^N2N3^ captures only ∼50% of Hb:Hp’s hemin (^max^ΔA_405_ ∼0.15) (**Fig. 5B**), whereas it completely removes the hemin from isolated metHb (^max^ΔA_405_ of ∼0.3) (**Fig. 5C**). Moreover, the N1 domain within IsdH^FL^ slows and limits hemin removal from isolated metHb, likely by competing with the extraction unit for αHb access, an effect not observed for Hb:Hp. Together, these findings suggest that an inherent feature of the Hb:Hp complex predisposes extraction unit binding toward βHb thereby limiting hemin removal from Hb:Hp.

**Figure 5:**
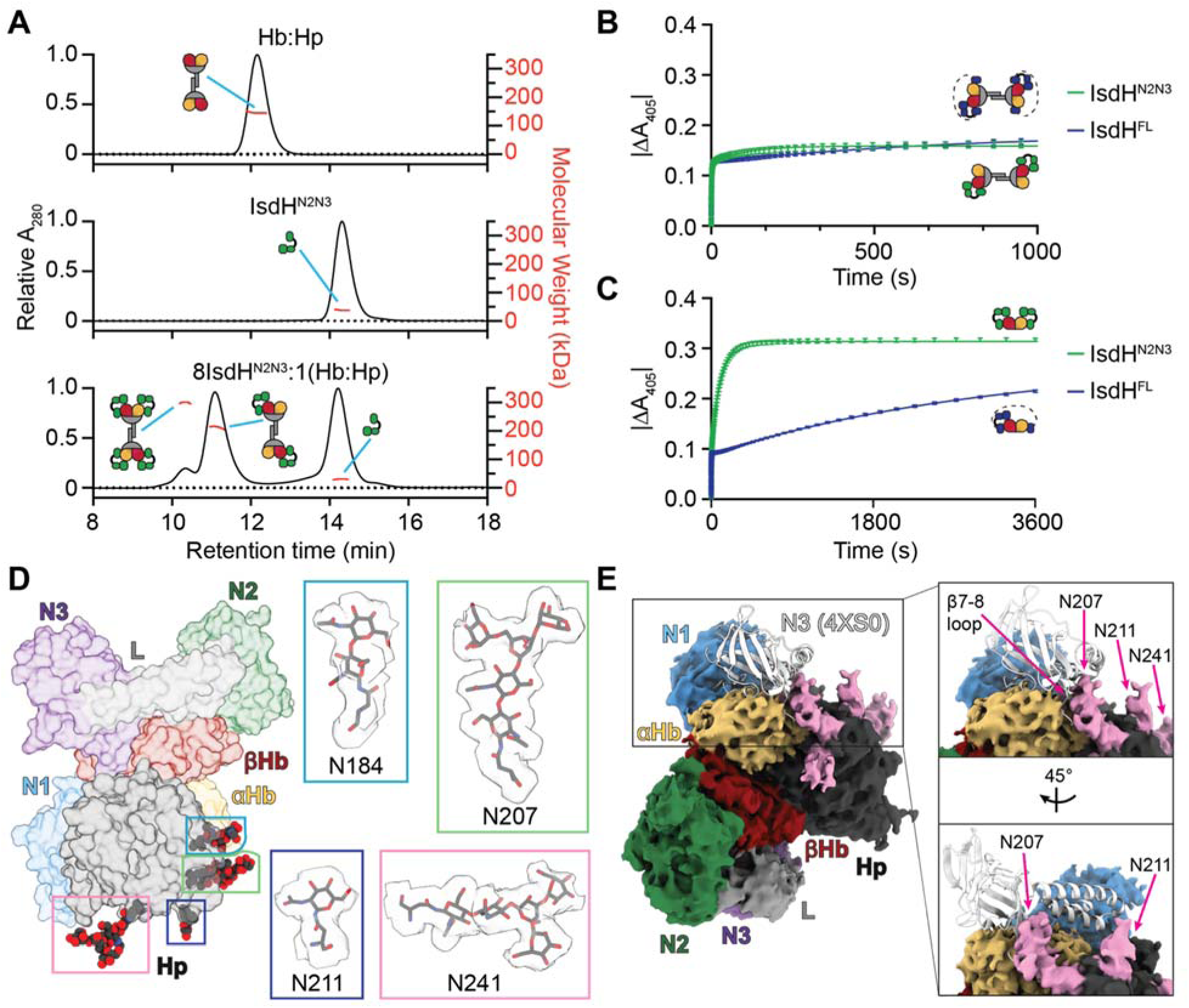
IsdH preferentially acquires βHb hemin from the Hb:Hp complex. (**A**) SEC-MALS of Hb:Hp (top), IsdH^N2N3^ (middle), and Hb:Hp + 8-fold excess IsdH^N2N3^ (bottom). Absorbance changes accompanying hemin capture by IsdH^N2N3^ (green) and IsdH^FL^ (blue) from (**B**) Hb:Hp and (**C**) Hb. (**D**) Each of Hp’s four glycosylation sites are visible in the cryo-EM map. (Left) Glycans on the model are shown as spheres colored by heteroatom and boxed in pink (N184), green (N207), dark blue (N211), and blue (N241). (Right) Glycans are shown expanded in boxes labelled with their corresponding Asn residue number on the right. The surrounding map region is depicted as transparent gray and contoured at 4 rmsd (threshold 0.025). (**E**) (Left) A crystal structure of an αHb-specific IsdH^N2N3^ mutant bound to Hb (white; PDB: 4XS0) (47) aligned to our cryo-EM structure approximates the extraction unit binding site on αHb within the Hb:Hp complex. The cryo-EM map contoured at 4 rmsd (threshold 0.025) was overlaid onto the aligned models. The map surrounding the glycans is colored pink, and the N3 β7-β8 loop and the glycans at N207, N211, and N241 are indicated with pink arrows (boxed on right).

Inspection of the cryo-EM map suggests that Hp glycosylation may limit extraction unit binding to αHb. Residues N184, N207, N211, and N241 in the Hp SP domain exhibit additional density attributable to sialylated biantennary or triantennary complex N-glycans (**Figs. 5D & S1A**) (77, 78). The map accommodates glycan structures ranging from a single GlcNAc residue to the conserved Man₃GlcNAc₂ pentasaccharide core; however, the outer antennae present at each site are unresolved, likely due to flexibility and microheterogeneity (79). Interestingly, the oligosaccharide at N207 on Hp is positioned to sterically hinder extraction unit binding to αHb. Alignment of an αHb-specific IsdH^N2N3^:Hb crystal structure (PDB: 4XS0) to the IsdH^FL^:Hb:Hp cryo-EM model and map reveals a clash between the N207-linked glycan and the N3 β7-β8 loop, which is critical for hemin extraction (**Fig. 5E**) (49). Notably, the N1 domain is smaller than the extraction unit, and its binding site on αHb is not obstructed by Hp glycosylation. Thus, Hp-associated glycans may bias hemin capture toward βHb by sterically restricting extraction unit access to αHb while permitting N1 binding.

## Discussion

*Staphylococcus aureus* acquires hemin from human Hb during infection, but Hb released from RBCs rapidly becomes inaccessible to bacteria as it is sequestered into Hb:Hp complexes and cleared by CD163-expressing macrophages (11, 13). We now present evidence that the *S. aureus* IsdH receptor is structurally adapted to overcome this barrier by selectively extracting hemin from βHb within Hb:Hp, and structural rationale for its impairment of CD163 binding, which is required for clearance. Biochemical measurements on full-length IsdH (IsdH^FL^) show its rapid removal of ∼50% of hemin from Hb:Hp, despite the complex’s high hemin affinity, and the post-extraction cryo-EM structure revels hemin is transferred from βHb to the receptor’s N2N3 extraction unit while the N1 domain is bound to αHb. We propose that βHb selectivity is driven by Hb:Hp glycosylation, not from intrinsic N2N3 specificity. In our cryo-EM map, N-linked glycans on the Hp serine protease domain are positioned to sterically occlude extraction-unit binding to αHb, while still permitting binding of the smaller N1 domain. This glycan-directed selectivity is supported by studies of the isolated extraction unit (IsdH^N2N3^), which, even when at high concentrations, predominantly forms a 2:1 complex with Hb:Hp, suggesting that only two of the four Hb chains are accessible for productive engagement. Moreover, although IsdH^N2N3^ can remove hemin from all globin chains in isolated metHb, it extracts only ∼50% of the hemin from Hb:Hp, mirroring the behavior of IsdH^FL^ and indicating that the Hb:Hp complex itself restricts extraction to half of the available sites.

The results reported here provide insight into the mechanism of hemin capture. While our structure captures the post-extraction complex, previously reported structures of the isolated extraction unit bound to HbCO (PDB: 9S3P) or to Hb:Hp^SP^ (an Hb dimer bound to recombinant, non-glycosylated Hp SP domain, PDB: 6TB2) likely represent pre- and mid-extraction intermediates, respectively (**Fig. 6**) (52, 62). Together, they support a plausible mechanism of transfer in which hemin migrates ∼13.4 Å from the βHb pocket to the receptor’s N3 domain. In the pre-transfer state, hemin remains bound to βHb, with its iron coordinated by proximal H92 and capped by distal H63. IsdH Y642 within N3 engages hemin’s 6-propionate, foreshadowing its eventual role as the axial ligand following extraction. As transfer proceeds, hemin moves toward the N3 domain, breaking its coordination bond with H92 and forming a new bond with distal H63, while nearby receptor residues undergo a disordered-to-ordered transition that creates a short 3_10_ helix that positions S563 for hydrogen bonding to the 7-propionate group. The reaction culminates in the post-transfer state visualized in our structure, in which βHb H63 is replaced by Y642 in the receptor as the axial ligand, while Y646 is positioned to hydrogen bond to Y642, as observed in other NEAT domains. During this process the hemin molecule rotates ∼95°, and H63 in βHb establishes new contacts to the 7-propionate. Transfer partially separates N3 and βHb, reducing their buried surface area by ∼300 Å^2^ and leaving the βHb’s empty pocket expanded and distorted (**Table S2**).

**Figure 6:**
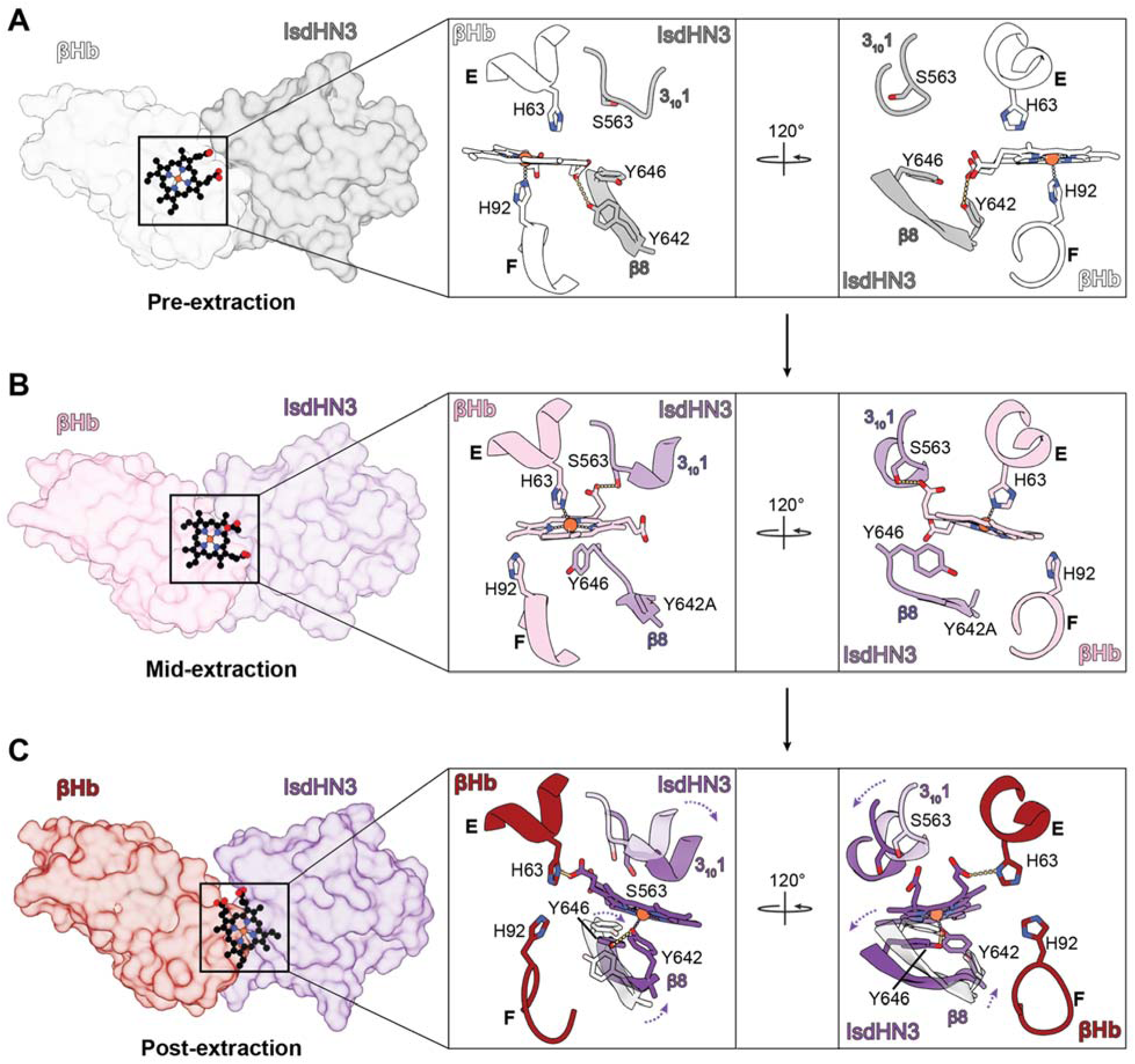
Snapshots of βHb hemin capture by IsdH. (**A**) Pre-extraction (IsdH^N2N3^ in complex with an HbCO tetramer; PDB: 9S3P), (**B**) mid-extraction (IsdH^N2N3^ harboring an inactivating Y642A mutation in complex with a presumed oxidized Hb dimer and recombinant Hp SP domain; PDB: 6TB2), and (**C**) the post-extraction IsdH^FL^:Hb:Hp cryo-EM structure reported herein. IsdH N3 and βHb are shown as transparent surfaces with hemin molecules shown in ball and stick representation (left). Additional protein chains are omitted for visual clarity. (Right) The heme/hemin environment is shown for each structure. The IsdH N3 α1 helix (mid-extraction: light purple, and post-extraction: dark purple) and β8 strand (pre-extraction: gray, and post extraction: dark purple) are shown aligned onto the post-transfer structure. N3 domain movement is indicated with dashed purple arrows.

Three-dimensional variability analysis of the IsdH^FL^:Hb:Hp cryo-EM data indicates that the receptor engages the βHb heme pocket through a dynamic interface in which the L domain’s α2 helix approaches and withdraws from the βHb F-helix. These motions resemble those proposed for hemin extraction from isolated Hb based on NMR and MD simulations, in which interdomain rearrangements transiently reposition the receptor’s L and N3 domains over the globin hemin pocket, juxtaposing two functional subsites on the receptor that are important for extraction (L-domain R492/Y495 and N3-domain V637/I640) (49, 74). Our cryo-EM data support a similar mechanism in Hb:Hp, as conformational heterogeneity is concentrated in the L and N3 domains and positions Y495 intermittently near the βHb F-helix, consistent with transient engagement of this L-domain subsite. Notably, this differs from previously observed behavior for HbCO-bound IsdH^N2N3^ (52), where motion is restricted and no heme is extracted, implying that the L-F-helix motions we observe are linked to productive capture. As our data indicate that IsdH accelerates hemin release from Hb:Hp to a similar extent as from Hb, these observations support a model in which IsdH transiently distorts the βHb F-helix to introduce strain that lowers the activation barrier for breaking the proximal H92-Fe coordination bond (46).

The N1 domain in IsdH emerges as a key determinant in prolonging microbial access to Hb by perturbing multiple pathways that normally clear it from circulation. IsdH interferes with CD163 recognition of the Hb:Hp complex on macrophages, thereby impairing receptor-mediated endocytosis (61, 62). Our results provide molecular insight into how the full-length receptor is optimized for this function, as comparison of our IsdH^FL^:Hb:Hp model with recently reported CD163:Hb:Hp structures suggests that simultaneous αHb and βHb engagement by IsdH sterically occludes distinct CD163 binding sites on Hb:Hp (**Fig. 7A-C**) (70–73). Furthermore, because N1 is connected to the extraction unit by a long disordered linker, the “cross-binding” between Hb:Hp complexes detected in our SEC-MALS and MS experiments suggests that IsdH can engage higher-order assemblies containing Hp2 phenotypes, thereby further disrupting CD163 binding. In addition to Hb:Hp, free Hb tetramers are also endocytosed via a secondary CD163-mediated clearance pathway (15). The ability of IsdH^FL^ to engage both globin subunits within Hb, as observed in our Hb:Hp-bound structure, is therefore also expected to impair this secondary CD163-mediated clearance pathway. Finally, a recent cryo-EM structure revealed a distinct mode of IsdH binding to Hb dimers, in which N1 engages a site that overlaps the Hp interface and may hinder the onset of Hp-mediated clearance (55). Collectively, these observations support a model in which IsdH disrupts host Hb clearance at multiple levels, ranging from interference with Hb:Hp complex formation to inhibition of CD163-mediated endocytosis of both free Hb and Hb:Hp complexes (**Fig. 7D**).

**Figure 7:**
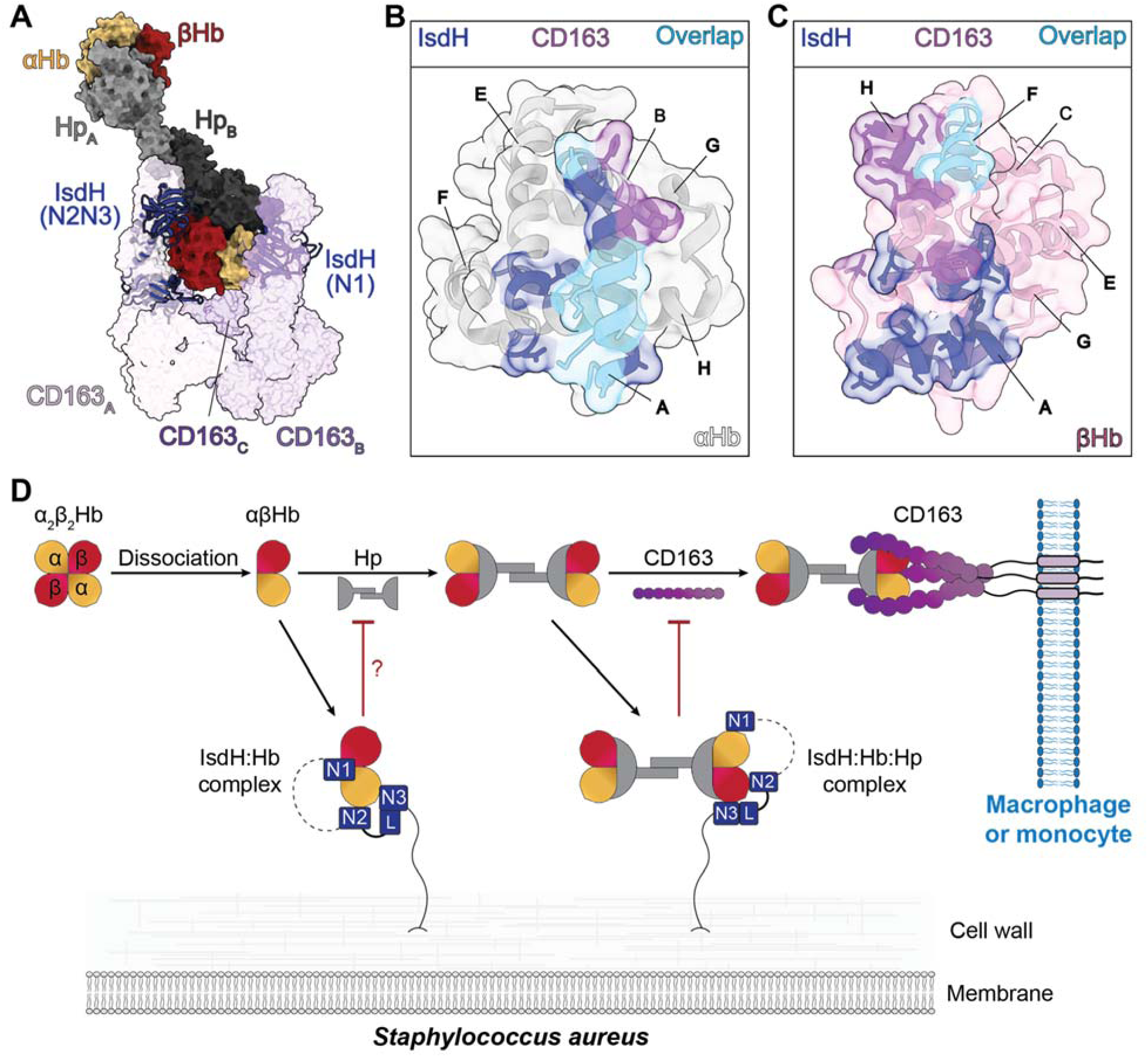
Overview of Hb clearance impedance by IsdH. (**A**) Alignment of the IsdH^FL^:Hb:Hp and CD163:Hb:Hp (PDB: 9NB6) complexes reveals that IsdH binding occludes two CD163 binding sites. Overlay of the IsdH (dark blue) and CD163 (purple) footprints on (**B**) αHb and (**C**) βHb. Shared contacts are shown in light blue. (**D**) Schematic of the primary human Hb clearance pathway and IsdH interference. Black arrows indicate known steps in clearance and in Hb or Hb:Hp capture by IsdH. Red capped lines indicate where clearance pathway component binding is known or thought to be impaired.

In conclusion, *S. aureus* may use its closely related IsdB and IsdH receptors to exploit distinct heme pools *in vivo*. IsdB appears dedicated to rapid hemin acquisition from free Hb, whereas IsdH, equipped with an additional N-terminal N1 binding domain, is specialized to remove hemin from βHb within Hb:Hp and to disrupt its clearance via CD163-mediated endocytosis. Paradoxically, however, IsdH appears optimized specifically for Hb:Hp, as our data show that the N1 domain actually diminishes its capacity to remove hemin from isolated Hb. To the best of our knowledge, *S. aureus* is unique among pathogens in encoding a receptor that actively extracts hemin from Hb:Hp, an abundant but notoriously recalcitrant source of heme iron. This capacity may reflect the ability of this lethal pathogen to colonize diverse host niches and underscores IsdH-mediated heme acquisition as an attractive target for anti-virulence therapies in antibiotic-resistant *S. aureus* infections.

## Materials and methods

### Protein production

Plasmids encoding IsdH^N2N3^ (residues 326-660) and H64V/V68F sperm whale myoglobin (apo-Mb^H64Y/V68F^) have been described previously (46, 80). The plasmid expressing IsdH^FL^ (residues 82-660) was constructed using standard methods using pET-28a as a scaffold and the IsdH sequence from *S. aureus* strain NCTC 10833. IsdH^FL^, IsdH^N2N3^, and apo-Mb were expressed and purified as previously described (74, 80, 81). Human HbA (Hb) was prepared from the blood of a healthy donor provided by the CFAR Virology Core Lab at the UCLA AIDS Institute, purified in the carbonmonoxy form and converted to metHb as previously described (74). Hb:Hp complexes were prepared by incubating 1 mg lyophilized Hp1-1, Hp2-1, or Hp2-2 (Athens Bioscience) with 5-fold molar excess Hb in 20 mM sodium phosphate, 150 mM NaCl pH 7.5 on ice for 2: 30 min. Excess Hb was removed using a Superdex S200 Increase 10/300 GL column (Cytiva). Hb:Hp complex-containing fractions were concentrated in 15 mL Amicon centrifugal filters (30 kDa MWCO), aliquoted, flash frozen in liquid N_2_, and stored at - 80°C.

### SEC-MALS and SEC-SAXS

Samples containing Hb:Hp, IsdH^FL^, and IsdH^N2N3^ were prepared at 3-5 mg/mL in 50 mM Tris-HCl, 150 mM NaCl, 1% glycerol, pH 7.5 (SAXS buffer) and submitted to the SIBYLS beamline (Advanced Light Source beamline 12.3.1, Lawrence Berkeley National Laboratory) (82). SEC-MALS and SEC-SAXS were performed using an Agilent 1290 high-pressure liquid chromatography (HPLC) system with a Shodex-KW 803 column equilibrated in SAXS buffer. SEC-MALS was conducted in-line using Wyatt Dawn Heleos multi-angle light scattering, DynaPro Titan quasi-elastic light scattering, and Optilab rEX refractometry detectors, and data were analyzed using Wyatt Astra 6 or 8 software packages. X-ray scattering data were collected continuously at a wavelength of 1.127 Å in 2 s frames on a Dectris PILATUS3 X 2M detector, with a sample to detector distance of 2 m. SEC-SAXS data were processed in ScÅtter IV (www.bioisis.net) and analyzed using BioXTAS RAW (83) and the ATSAS Suite (84). DENSS was used to reconstruct 20 density maps, followed by averaging and refinement (63).

### EM sample preparation and data acquisition

Negative stain electron micrographs were prepared by depositing 3 µL of solution containing 12.5 µg/mL Hb:Hp or Hb:Hp with 4-fold excess apo-IsdH^FL^ onto 300 mesh formvar/carbon grids. Grids were washed once with nanopure water and stained for 2 minutes with 1% uranyl acetate prior to imaging using an FEI Tecnai 12 electron microscope.

To form the complex for cryo-EM studies, 1 µM apo-IsdH^FL^ and 1.25 µM Hb:Hp1-1 complex were combined in 50 mM Tris-HCl, 150 mM NaCl, and 0.25% glycerol and incubated on ice for 30 min. 3 µL of solution was applied to glow-discharged (PELCO easiGlow, Ted Pella) Quantifoil R 1.2/1.3, 300 mesh, Cu grids (Electron Microscopy Sciences) in a Vitrobot Mark IV (Thermo Fisher Scientific) with the chamber set to 100% relative humidity and 4°C. Grids were blotted with blot force −6, blot time 12 or 14 s, drain time 1 s and plunge frozen in liquid ethane. Cryo-EM data acquisition was performed using a Titan Krios G4 (Thermo Fisher Scientific) equipped with a Selectris X energy filter and a Falcon 4i Direct Electron Detector. Movies were collected with a total fluence of 50 e^−^/Å^2^ at 130,000x magnification (pixel size 0.97 Å) with a target defocus range of −1.0 to −2.5 µM. 14,847 movies were collected across two datasets using EPU software (Thermo Fisher Scientific). Data collection parameters are summarized in **Table S3**.

### Image processing and model building

All cryo-EM data processing was conducted in CryoSPARC v4.6.2-v4.7.0 (Structura Biotechnology) as depicted in (**Fig. S9**) and detailed in the supplementary materials and methods. An initial model of the IsdH^FL^:Hb:Hp complex was generated using AlphaFold 3 (85), docked into the cryo-EM map in ChimeraX (86), and relaxed into the unsharpened map using Rosetta (76). Iterative rounds of manual model adjustment into the unsharpened cryo-EM map and refinement were conducted in Coot and Phenix, respectively (87, 88). ChimeraX was used for model and map visualization and for rendering figures. Cavity volume calculations were performed using pyKVFinder (89).

### Hemin transfer experiments

Spectral changes occurring upon mixing 4 µM apo-IsdH^FL^ and 5 µM Hb:Hp1-1 (all Hb:Hp concentrations are reported on a hemin basis for kinetic experiments) with Hb in the met or carbonmonoxy form were initially monitored using a Shimadzu 1800 UV spectrophotometer for 20 min in 20 mM sodium phosphate, 150 mM NaCl, 450 mM sucrose pH 7.5 (heme transfer buffer). The rapid spectral change observed when metHb:Hp was used as the heme source prompted use of an Applied Photophysics SX-20 stopped-flow spectrophotometer to better resolve time-dependent spectral changes accompanying heme capture. 5 µM metHb:Hp^mix^ was mixed with 25 µM IsdH, and spectral changes were monitored for 15 min at 25°C using a photodiode array detector.

Hemin capture kinetics were monitored using an Applied Photophysics SX-20 stopped-flow spectrophotometer in triplicate and were performed and analyzed as previously described (74). Briefly, 5 µM metHb:Hp^mix^ was rapidly mixed with 1-150 µM apo-IsdH^FL^ or 150 µM apo-Mb in heme transfer buffer, and absorbance changes at 405 nm were monitored for 1 hour at 25°C using an absorbance photomultiplier tube detector. Additional experiments were conducted using 5 µM metHb, metHb:Hp1-1, or metHb:Hp2-2 as the hemin source and 150 µM apo-IsdH^FL^, apo-IsdH^N2N3^, or apo-Mb as the hemin acceptor. Resulting curves were fit to a biphasic exponential decay equation using GraphPad Prism version 10.2.1.

### Kinetics of passive Hb- and Hb:Hp-hemin dissociation

A passive Hb:Hp hemin release assay using apo-Mb as the hemin acceptor was conducted as described previously (90). Briefly, 50 µM apo-Mb was combined with 0, 2.5, 5, 10, 25, or 50 µM Hp1-1 in heme transfer buffer in a 384-well plate (Greiner Bio-One). Reactions were initiated by injection of metHb to a final concentration of 5 µM in a SpectraMax iD3 plate reader (Molecular Devices). The absorbance change at 405 nm was monitored at 25°C every 5 min for 20h. Additional experiments were performed using 5 µM pre-complexed Hb:Hp1-1, Hb:Hp^mix^, and Hb:Hp2-2 and were initiated by injection of 50 µM apo-Mb. Experiments were performed in triplicate and the absorbance traces were plotted and fit to a two-phase exponential equation as described above.

## Acknowledgements

This research was funded by grants from the National Institutes of Health R01 AI161828 (RTC), NIH-NIGMS R35GM128867 (JAR), and R35GM145286 (JAL) and was also performed as part of STROBE, an NSF Science and Technology Center through Grant DMR-1548924. J.A.R. was further supported as a Packard Fellow. J.S. was partially supported by a NIH NIGMS-funded predoctoral fellowship (T32 GM136614). A.K.G. acknowledges support from the NIH National Institute of Dental & Craniofacial Research (T90DE030860). We acknowledge the use of instruments at the Electron Imaging Center for Nanomachines supported by UCLA and grants from the NIH (1S10OD018111). SAXS data were collected at SIBYLS which is supported by the DOE-BER IDAT DE-AC02-05CH11231 and NIGMS ALS-ENABLE (P30 GM124169 and S10OD018483). We thank members of the Clubb and Rodriguez labs for useful discussions and advice and David Strugatsky for his assistance in cryo-EM data collection.

## References

1. J. P. Steinberg, C. C. Clark, B. O. Hackman, Nosocomial and Community-Acquired Staphylococcus aureus Bacteremias from 1980 to 1993: Impact of Intravascular Devices and Methicillin Resistance. Clin. Infect. Dis. 23, 255–259 (1996).

2. A. S. Lee, et al., Methicillin-resistant Staphylococcus aureus. Nat. Rev. Dis. Primers 4, 18033 (2018).

3. Centers for Disease Control and Prevention (U.S.), “Antibiotic resistance threats in the United States, 2019” (Centers for Disease Control and Prevention (U.S.), 2019).

4. C. J. L. Murray, et al., Global burden of bacterial antimicrobial resistance in 2019: a systematic analysis. Lancet 399, 629–655 (2022).

5. United Nations Environment Programme, Bracing for Superbugs: Strengthening Environmental Action in the One Health Response to Antimicrobial Resistance (United Nations, 2023).

6. C. C. Murdoch, E. P. Skaar, Nutritional immunity: the battle for nutrient metals at the host–pathogen interface. Nat. Rev. Microbiol. 20, 657–670 (2022).

7. Q. Ru, et al., Iron homeostasis and ferroptosis in human diseases: mechanisms and therapeutic prospects. Sig. Transduct. Target Ther. 9, 271 (2024).

8. E. P. Skaar, Iron-Source Preference of Staphylococcus aureus Infections. Science 305, 1626–1628 (2004).

9. J. E. Cassat, E. P. Skaar, Iron in Infection and Immunity. Cell Host Microbe 13, 509–519 (2013).

10. J. E. Choby, E. P. Skaar, Heme Synthesis and Acquisition in Bacterial Pathogens. J. Mol. Biol. 428, 3408–3428 (2016).

11. M. Kristiansen, J. H. Graversen, C. Jacobsen, O. Sonne, S. K. Moestrup, Identification of the haemoglobin scavenger receptor. Nature 409, 198–201 (2001).

12. C. B. F. Andersen, et al., Haptoglobin. Antioxid. Redox Signal. 26, 814–831 (2017).

13. A. di Masi, et al., Haptoglobin: From hemoglobin scavenging to human health. Mol. Asp. Med. 73, 100851 (2020).

14. D. J. Schaer, et al., CD163 is the macrophage scavenger receptor for native and chemically modified hemoglobins in the absence of haptoglobin. Blood 107, 373–380 (2006).

15. R. X. Zhou, M. K. Higgins, Structural basis for hemoglobin scavenging by CD163 reveals mechanism of ligand promiscuity. PLoS Biol. 24, e3003788 (2026).

16. Z. Hrkal, Z. Vodrážka, I. Kalousek, Transfer of Heme from Ferrihemoglobin and Ferrihemoglobin Isolated Chains to Hemopexin. Eur. J. Biochem. 43, 73–78 (1974).

17. V. Hvidberg, et al., Identification of the receptor scavenging hemopexin-heme complexes. Blood 106, 2572–2579 (2005).

18. S. K. Mazmanian, et al., Passage of Heme-Iron Across the Envelope of Staphylococcus aureus. Science 299, 906–909 (2003).

19. A. Dryla, D. Gelbmann, A. Von Gabain, E. Nagy, Identification of a novel iron regulated staphylococcal surface protein with haptoglobin-haemoglobin binding activity: Staphylococcal haptoglobin receptor. Mol. Microbiol. 49, 37–53 (2003).

20. V. J. Torres, G. Pishchany, M. Humayun, O. Schneewind, E. P. Skaar, *Staphylococcus aureus* IsdB Is a Hemoglobin Receptor Required for Heme Iron Utilization. J. Bacteriol. 188, 8421–8429 (2006).

21. K. Krishna Kumar, et al., Structural Basis for Hemoglobin Capture by Staphylococcus aureus Cell-surface Protein, IsdH. J. Biol. Chem. 286, 38439–38447 (2011).

22. H. Zhu, et al., Pathway for Heme Uptake from Human Methemoglobin by the Iron-regulated Surface Determinants System of *Staphylococcus aureus*. J. Biol. Chem. 283, 18450–18460 (2008).

23. T. Spirig, et al., Staphylococcus aureus Uses a Novel Multidomain Receptor to Break Apart Human Hemoglobin and Steal Its Heme. J. Biol. Chem. 288, 1065–1078 (2013).

24. S. K. Mazmanian, H. Ton-That, K. Su, O. Schneewind, An iron-regulated sortase anchors a class of surface protein during *Staphylococcus aureus* pathogenesis. Proc. Natl. Acad. Sci. U.S.A. 99, 2293–2298 (2002).

25. N. Muryoi, et al., Demonstration of the Iron-regulated Surface Determinant (Isd) Heme Transfer Pathway in Staphylococcus aureus. J. Biol. Chem. 283, 28125–28136 (2008).

26. L. A. Adolf, et al., Functional membrane microdomains and the hydroxamate siderophore transporter ATPase FhuC govern Isd-dependent heme acquisition in Staphylococcus aureus. eLife 12, e85304 (2023).

27. E. P. Skaar, A. H. Gaspar, O. Schneewind, IsdG and IsdI, Heme-degrading Enzymes in the Cytoplasm of Staphylococcus aureus. J. Biol. Chem. 279, 436–443 (2004).

28. M. A. Andrade, F. D. Ciccarelli, C. Perez-Iratxeta, P. Bork, NEAT: a domain duplicated in genes near the components of a putative Fe3+siderophore transporter from Gram-positive pathogenic bacteria. Genome Biol. 3, research0047.1 (2002).

29. J. C. Grigg, G. Ukpabi, C. F. M. Gaudin, M. E. P. Murphy, Structural biology of heme binding in the Staphylococcus aureus Isd system. J. Inorg. Biochem. 104, 341–348 (2010).

30. K. Ellis-Guardiola, B. J. Mahoney, R. T. Clubb, NEAr Transporter (NEAT) Domains: Unique Surface Displayed Heme Chaperones That Enable Gram-Positive Bacteria to Capture Heme-Iron From Hemoglobin. Front. Microbiol. 11, 607679 (2021).

31. A. Dryla, et al., High-Affinity Binding of the Staphylococcal HarA Protein to Haptoglobin and Hemoglobin Involves a Domain with an Antiparallel Eight-Stranded β-Barrel Fold. J. Bacteriol. 189, 254–264 (2007).

32. J. C. Grigg, C. L. Vermeiren, D. E. Heinrichs, M. E. P. Murphy, Heme Coordination by *Staphylococcus aureus* IsdE. J. Biol. Chem. 282, 28815–28822 (2007).

33. J. C. Grigg, C. L. Vermeiren, D. E. Heinrichs, M. E. P. Murphy, Haem recognition by a Staphylococcus aureus NEAT domain. Mol. Microbiol. 63, 139–149 (2007).

34. R. M. Pilpa, et al., Functionally Distinct NEAT (NEAr Transporter) Domains within the Staphylococcus aureus IsdH/HarA Protein Extract Heme from Methemoglobin. J. Biol. Chem. 284, 1166–1176 (2009).

35. M. Watanabe, et al., Structural Basis for Multimeric Heme Complexation through a Specific Protein-Heme Interaction. J. Biol. Chem. 283, 28649–28659 (2008).

36. C. F. M. Gaudin, J. C. Grigg, A. L. Arrieta, M. E. P. Murphy, Unique Heme-Iron Coordination by the Hemoglobin Receptor IsdB of *Staphylococcus aureus*. Biochemistry 50, 5443–5452 (2011).

37. R. Macdonald, B. J. Mahoney, K. Ellis-Guardiola, A. Maresso, R. T. Clubb, NMR experiments redefine the hemoglobin binding properties of bacterial NEAr-iron Transporter domains. Protein Sci. 28, 1513–1523 (2019).

38. D. Akinbosede, R. Chizea, S. A. Hare, Pirates of the haemoglobin. Microb. Cell 9, 84–102 (2022).

39. J. R. Sheldon, H. A. Laakso, D. E. Heinrichs, Iron Acquisition Strategies of Bacterial Pathogens. Microbiol. Spectr. 4, 4.2.05 (2016).

40. J. R. Sheldon, D. E. Heinrichs, Recent developments in understanding the iron acquisition strategies of gram positive pathogens. FEMS Microbiol. Rev. 39, 592–630 (2015).

41. H. Contreras, N. Chim, A. Credali, C. W. Goulding, Heme uptake in bacterial pathogens. Curr. Opin. Chem. Biol. 19, 34–41 (2014).

42. C. L. Nobles, A. W. Maresso, The theft of host heme by Gram-positive pathogenic bacteria. Metallomics 3, 788 (2011).

43. M. I. Hood, E. P. Skaar, Nutritional immunity: transition metals at the pathogen–host interface. Nat. Rev. Microbiol. 10, 525–537 (2012).

44. M. Sjodt, et al., The PRE-Derived NMR Model of the 38.8-kDa Tri-Domain IsdH Protein from Staphylococcus aureus Suggests That It Adaptively Recognizes Human Hemoglobin. J. Mol. Biol. 428, 1107–1129 (2016).

45. H. Zhu, D. Li, M. Liu, V. Copié, B. Lei, Non-Heme-Binding Domains and Segments of the Staphylococcus aureus IsdB Protein Critically Contribute to the Kinetics and Equilibrium of Heme Acquisition from Methemoglobin. PLoS ONE 9, e100744 (2014).

46. M. Sjodt, et al., Energetics underlying hemin extraction from human hemoglobin by *Staphylococcus aureus*. J. Biol. Chem. 293, 6942–6957 (2018).

47. C. F. Dickson, D. A. Jacques, R. T. Clubb, J. M. Guss, D. A. Gell, The structure of haemoglobin bound to the haemoglobin receptor IsdH from *Staphylococcus aureus* shows disruption of the native α-globin haem pocket. Acta Crystallogr. D Biol. Crystallogr. 71, 1295–1306 (2015).

48. C. F. M. Bowden, et al., Structure–function analyses reveal key features in Staphylococcus aureus IsdB-associated unfolding of the heme-binding pocket of human hemoglobin. J. Biol. Chem. 293, 177–190 (2018).

49. K. Ellis-Guardiola, et al., The Staphylococcus aureus IsdH Receptor Forms a Dynamic Complex with Human Hemoglobin that Triggers Heme Release via Two Distinct Hot Spots. J. Mol. Biol. 432, 1064–1082 (2020).

50. O. De Bei, et al., Cryo-EM structures of staphylococcal IsdB bound to human hemoglobin reveal the process of heme extraction. Proc. Natl. Acad. Sci. U.S.A. 119, e2116708119 (2022).

51. O. De Bei, et al., Time-resolved X-ray solution scattering unveils the events leading to hemoglobin heme capture by staphylococcal IsdB. Nat. Commun. 16, 1361 (2025).

52. V. B. Comani, et al., Hemoglobin receptor redundancy in Staphylococcus aureus: molecular flexibility as a determinant of divergent hemophore activity. J. Struct. Biol. X 12, 100138 (2025).

53. C. F. M. Bowden, M. M. Verstraete, L. D. Eltis, M. E. P. Murphy, Hemoglobin Binding and Catalytic Heme Extraction by IsdB Near Iron Transporter Domains. Biochemistry 53, 2286–2294 (2014).

54. B. A. Fonner, et al., Solution Structure and Molecular Determinants of Hemoglobin Binding of the First NEAT Domain of IsdB in *Staphylococcus aureus*. Biochemistry 53, 3922–3933 (2014).

55. V. B. Comani, et al., Refining the mechanism of heme acquisition from free hemoglobin by Staphylococcus aureus IsdH. Proc. Natl. Acad. Sci. U.S.A. 123, e2601134123 (2026).

56. H. K. Kim, et al., IsdA and IsdB antibodies protect mice against Staphylococcus aureus abscess formation and lethal challenge. Vaccine 28, 6382–6392 (2010).

57. G. Pishchany, et al., Specificity for Human Hemoglobin Enhances Staphylococcus aureus Infection. Cell Host Microbe 8, 544–550 (2010).

58. F. Vallelian, P. W. Buehler, D. J. Schaer, Hemolysis, free hemoglobin toxicity, and scavenger protein therapeutics. Blood 140, 1837–1844 (2022).

59. R. Larsen, Z. Gouveia, M. P. Soares, R. Gozzelino, Heme cytotoxicity and the pathogenesis of immune-mediated inflammatory diseases. Front. Pharmacol. 3, 77 (2012).

60. D. J. Schaer, F. Vinchi, G. Ingoglia, E. Tolosano, P. W. Buehler, Haptoglobin, hemopexin, and related defense pathways - basic science, clinical perspectives, and drug development. Front. Physiol. 5, 415 (2014).

61. K. L. Sæderup, et al., The Staphylococcus aureus Protein IsdH Inhibits Host Hemoglobin Scavenging to Promote Heme Acquisition by the Pathogen. J. Biol. Chem. 291, 23989–23998 (2016).

62. J. H. Mikkelsen, K. Runager, C. B. F. Andersen, The human protein haptoglobin inhibits IsdH-mediated heme-sequestering by Staphylococcus aureus. J. Biol. Chem. 295, 1781–1791 (2020).

63. T. D. Grant, Ab initio electron density determination directly from solution scattering data. Nat. Methods 15, 191–193 (2018).

64. K. Stødkilde, M. Torvund-Jensen, S. K. Moestrup, C. B. F. Andersen, Structural basis for trypanosomal haem acquisition and susceptibility to the host innate immune system. Nat. Commun. 5, 5487 (2014).

65. H. F. Bunn, J. H. Jandl, Exchange of Heme among Hemoglobins and between Hemoglobin and Albumin. J. Biol. Chem. 243, 465–475 (1968).

66. M. Lipiski, et al., Human Hp1-1 and Hp2-2 Phenotype-Specific Haptoglobin Therapeutics Are Both Effective *In Vitro* and in Guinea Pigs to Attenuate Hemoglobin Toxicity. Antioxid. Redox Signal. 19, 1619–1633 (2013).

67. T. L. Mollan, et al., Redox properties of human hemoglobin in complex with fractionated dimeric and polymeric human haptoglobin. Free Radic. Biol. Med. 69, 265–277 (2014).

68. C. B. F. Andersen, et al., Structure of the haptoglobin–haemoglobin complex. Nature 489, 456–459 (2012).

69. M. Przybylska, H. M. Sheppard, S. Szilágyi, Crystallization of the haptoglobin–hemoglobin complex. Acta Crystallogr. D Biol. Crystallogr. 55, 883–884 (1999).

70. A. Etzerodt, et al., The Cryo-EM structure of human CD163 bound to haptoglobin-hemoglobin reveals molecular mechanisms of hemoglobin scavenging. Nat. Commun. 15, 10871 (2024).

71. R. X. Zhou, M. K. Higgins, Scavenger receptor CD163 multimerises to allow uptake of diverse ligands. Nat. Commun. 16, 6623 (2025).

72. C.-S. Huang, et al., Structural elucidation of the haptoglobin–hemoglobin clearance mechanism by macrophage scavenger receptor CD163. PLoS Biol. 23, e3003264 (2025).

73. H. Xu, X. Song, X. Su, Calcium-dependent oligomerization of scavenger receptor CD163 facilitates the endocytosis of ligands. Nat. Commun. 16, 6679 (2025).

74. J. Clayton, et al., Directed Inter-domain Motions Enable the IsdH Staphylococcus aureus Receptor to Rapidly Extract Heme from Human Hemoglobin. J. Mol. Biol. 434, 167623 (2022).

75. A. Punjani, D. J. Fleet, 3D variability analysis: Resolving continuous flexibility and discrete heterogeneity from single particle cryo-EM. J. Struct. Biol. 213, 107702 (2021).

76. M. D. Tyka, et al., Alternate States of Proteins Revealed by Detailed Energy Landscape Mapping. J. Mol. Biol. 405, 607–618 (2011).

77. P. Pompach, et al., Site-specific Glycoforms of Haptoglobin in Liver Cirrhosis and Hepatocellular Carcinoma. Mol. Cell. Proteomics 12, 1281–1293 (2013).

78. S. Zhang, K. Jiang, C. Sun, H. Lu, Y. Liu, Quantitative analysis of site-specific N-glycans on sera haptoglobin beta-chain in liver diseases. ABBS 45, 1021–1029 (2013).

79. S. Tamara, V. Franc, A. J. R. Heck, A wealth of genotype-specific proteoforms fine-tunes hemoglobin scavenging by haptoglobin. Proc. Natl. Acad. Sci. U.S.A. 117, 15554–15564 (2020).

80. M. S. Hargrove, et al., His64(E7)–>Tyr apomyoglobin as a reagent for measuring rates of hemin dissociation. J. Biol. Chem. 269, 4207–4214 (1994).

81. K. Ellis-Guardiola, J. Soule, R. Clubb, Methods for the Extraction of Heme Prosthetic Groups from Hemoproteins. Bio Protoc. 11 (2021).

82. S. Classen, et al., Implementation and performance of SIBYLS: a dual endstation small-angle X-ray scattering and macromolecular crystallography beamline at the Advanced Light Source. J. Appl. Crystallogr. 46, 1–13 (2013).

83. J. B. Hopkins, BioXTAS RAW 2: new developments for a free open-source program for small-angle scattering data reduction and analysis. J. Appl. Crystallogr. 57, 194–208 (2024).

84. K. Manalastas-Cantos, et al., *ATSAS 3.0* : expanded functionality and new tools for small-angle scattering data analysis. J. Appl. Crystallogr. 54, 343–355 (2021).

85. J. Abramson, et al., Accurate structure prediction of biomolecular interactions with AlphaFold 3. Nature 630, 493–500 (2024).

86. E. C. Meng, et al., UCSF ChimeraX: Tools for structure building and analysis. Protein Sci. 32, e4792 (2023).

87. P. Emsley, K. Cowtan, *Coot* : model-building tools for molecular graphics. Acta Crystallogr. D Biol. Crystallogr. 60, 2126–2132 (2004).

88. P. V. Afonine, et al., Real-space refinement in *PHENIX* for cryo-EM and crystallography. Acta Crystallogr. D Struct. Biol. 74, 531–544 (2018).

89. J. V. D. S. Guerra, et al., pyKVFinder: an efficient and integrable Python package for biomolecular cavity detection and characterization in data science. BMC Bioinformatics 22, 607 (2021).

90. R. Macdonald, et al., The Shr receptor from *Streptococcus pyogenes* uses a cap and release mechanism to acquire heme–iron from human hemoglobin. Proc. Natl. Acad. Sci. U.S.A. 120, e2211939120 (2023).

