## Supplemental Material for "Cryo-EM provides insight into how the *Staphylococcus aureus* IsdH receptor removes hemin from the hemoglobin:haptoglobin complex"

**This file includes:**

Supplementary Materials and Methods

Figures S1 to S10

Tables S1 to S3

Supplementary References

**Supplementary Materials and Methods**

**Cryo-EM processing and 3D Variability Analysis.** 14,847 movies underwent patch motion correction and patch CTF estimation. After micrograph curation, 13,378 exposures were retained. Particles were initially selected from 50% of the exposures using a pre-trained Topaz model (ResNet16, 64 units) (1). Particles were classified in 2D, and projections displaying clear secondary structure features were used to train a new Topaz model. Extraction with that Topaz model produced 457,405 particles which were subjected to 2D classification and *ab-initio* reconstruction into 3 volumes corresponding to the fully occupied IsdH^FL^:Hb:Hp complex and two partially occupied species. Particles assigned to the fully occupied complex underwent two rounds of heterogeneous refinement using the three *ab-initio* volumes and two decoy volumes generated from junk 2D classes. Refinement revealed map anisotropy consistent with preferred orientation (**Fig. S10**), prompting training of a second Topaz model using only rare views.

In parallel, particles were extracted from all micrographs using both Topaz models. After independent cleanup by two rounds of heterogeneous refinement, 371,043 and 398,565 particles were retained from the rare-view and initial Topaz models, respectively. Particles from the rare-view-trained model underwent non-uniform refinement and 3D classification to identify better-resolved subvolumes with improved orientation diagnostics. The three most populated 3D classes were used as reference volumes for heterogeneous refinement using the combined particle stack (659,051 particles). Subsequent rounds of non-uniform refinement, 3D classification, and curation by alignment error, CTF refinement, and reference-based motion correction further improved the map. The final consensus reconstruction was produced by subsetting the particle stack based on per particle scale, homogeneous refinement, and a final round of non-uniform refinement using a mask encompassing IsdH N1, IsdH N2N3, an Hb dimer, and Hp SP.

The final particle stack containing 62,345 images was subjected to 3D variability analysis using an 8 Å filter resolution (2). Results were visualized in intermediates mode with 10 clusters. The central 6 particle clusters produced maps suitable for further analysis, and the IsdHFL:Hb:Hp model was fit to these using Rosetta FastRelax (3).

**Native mass spectrometry.** Proteins for native MS experiments were buffer exchanged into 200 mM ammonium acetate (pH 6.9) using two successive rounds of Zeba spin columns (7 kDa MWCO, 0.5 mL column; Thermo Fisher Scientific), each pre-equilibrated by 10 successive flushes with the same buffer. IsdH was incubated with Hb:Hp complexes at 1 µM and 1.25 µM respectively, prior to ionization. Ionization was performed using native nano-electrospray ionization (nESI) with platinum-coated borosilicate emitters on a Thermo Q-Exactive Ultra High Mass Range (UHMR) mass spectrometer modified with an e-MSion ExD cell in place of the ion transfer multipole. A spray voltage of 0.7 to 1.1 kV was applied, with an S/N threshold of 0 and resolution of 3,125 at 400 m/z. Haptoglobin’s glycoforms caused complex and heterogeneous peak overlap in m/z space, so masses of Hp-containing complexoforms were obtained using electron-capture charge reduction (ECCR) to resolve overlapping charge states. Voltage settings for ExD cell lenses (L), lens magnets (LM), and filament bias (FB) used for ECCR were adapted from Le Huray and colleagues (L1 & L7 = 0 V, L2 & L6 = -25 V, LM3 & LM5 = 7 V, L4 = 7 V, FB = 1 V) (4). An in-source trapping voltage of -20 V, trapping gas pressure setting of 8, and an HCD cell energy of 1 V were applied. Ion transfer target was set to high m/z and detector m/z optimization was set to low m/z. Spectra were acquired over a scan range of 1,000 to 25,000 m/z. For each sample analyzed by native MS, multiple ECCR spectra were collected from consecutive isolation windows spanning 4,400 to 11,700 m/z (100 m/z spacing, 2 m/z overlap) for 6 microscans each with a 500 ms injection time. Composite sum zero charge mass spectra were generated following deconvolution of ECCR spectra in UniDec (5).

**NMR spectroscopy.** A sample of uniformly ^15^N-labeled IsdH^FL^ was prepared by expression in minimal media supplemented with ^15^NH_4_Cl (6), with purification of resulting protein performed as above. The final sample was prepared at a concentration of 100 µM in NMR buffer composed of 50 mM sodium phosphate, 100 mM NaCl, 8% D_2_O, pH 6.5. 2-D ^15^N-^1^H TROSY-HSQC spectra were acquired at 298 K on a Bruker AVANCE NEO 800 MHz (18.8 T) spectrometer equipped with a triple resonance cryogenic probe (7).

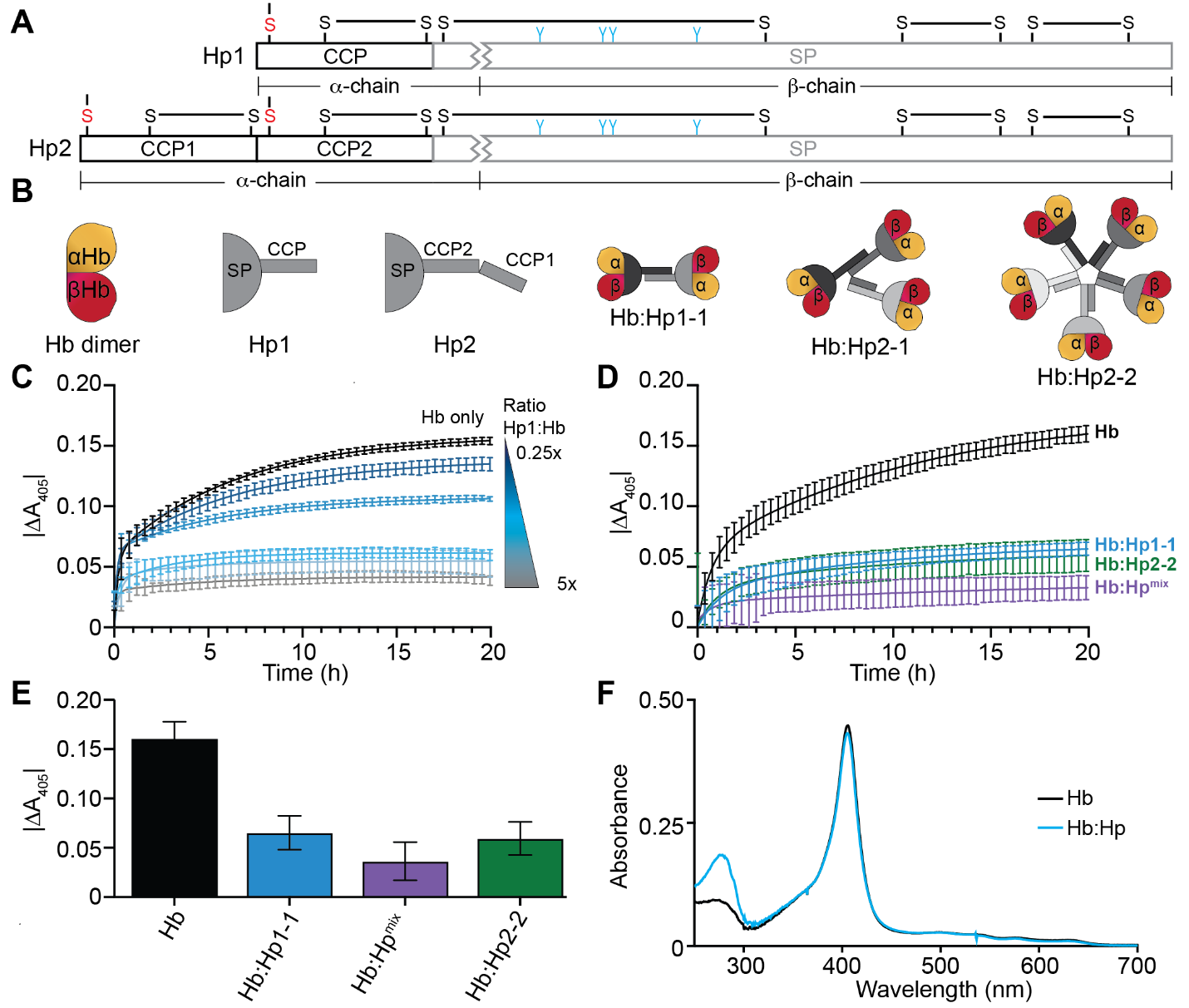

**Figure S1: Hp binding limits hemin release from metHb.**

(**A**) Domain schematic for the Hp1 and Hp2 variants of Hp (8, 9). Both proteins undergo cleavage N-terminal to Ile162 (Hp2 numbering) yielding an αHp chain containing the CCP domain(s) (black) and 13 residues of the SP domain (gray), and a βHp chain containing the remainder of the SP domain (gray). These cleavage products are linked by a disulfide bond, and also contain additional intrachain disulfide bonds. In the figure, cysteine residues connecting the CCP domains in distinct chains are shown as red “S” letters and N-linked glycosylation sites in the SP domains are shown as blue branched structures. (**B**) Cartoon representations of an Hb dimer, Hp1, Hp2, and Hb:Hp complexes. Hp1 homozygotes produce Hb:Hp1-1 dimers, whereas Hp2-harboring humans produce higher order oligomers. Notably, the Hp2-1 phenotype results in Hp1-1 dimers, Hp2-2 oligomers, and oligomeric species containing both Hp1 and Hp2. (**C**) Hemin release measurements from metHb in the presence of Hp1-1. The graph shows the time-dependent changes in Soret region (405 nm) after mixing Hb or the Hb:Hp complex with excess apo-Mb (sperm whale apo-myoglobin containing H64V and V68F mutations), an established hemin scavenging reagent which captures hemin that is spontaneously released from Hb or Hb:Hp (10). The reactions contain 50 µM apo-Mb and 5 µM metHb in the presence of 0, 0.25, 0.5, 1, 2.5, or 5x Hp1-1. A control experiment (black curve) in which isolated metHb is incubated with apo-Mb show expected bi-phasic absorbance changes described by *k*_fast_ and *k*_slow_ rate constants that report on hemin egress from the β- and α-globins, respectively (1.43 x 10^-3^ ± 0.10 x 10^-3^ s⁻¹ and 4.24 x 10^-5^ ± 0.09 x 10^-5^ s⁻¹, respectively). The addition of Hp1 impairs hemin release from metHb in a concentration-dependent manner. Error bars represent the standard deviation of three replicates. (**D**) Absorbance changes at 405 nm upon mixing 5 µM Hb, pre-complexed Hb:Hp1-1, Hb:Hp^mix^ (pooled Hp from blood containing the 1-1, 2-1, and 2-2 phenotypes), and Hb:Hp2-2 with 10-fold excess apo-Mb were tracked for 20 hours. Error bars represent the standard deviation of three replicates. (**E**) The total absorbance changes for Hb and the Hb:Hp complexes over the 20 hour time-course indicate that Hp limits hemin loss from Hb in a phenotype-independent manner. Error bars represent the standard error of the calculated |∆A_405_|. (**F**) A UV-Vis spectral overlay of 2.5 µM metHb and metHb:Hp1-1. Hb and Hb:Hp hemin concentrations were determined by the hemochome assay prior to acquisition of UV-Vis spectra. The metHb and metHb:Hp spectra do not differ appreciably (∆A_405_ ~ 0.01), thus Hp binding does not account for the spectral changes observed in our hemin transfer assays.

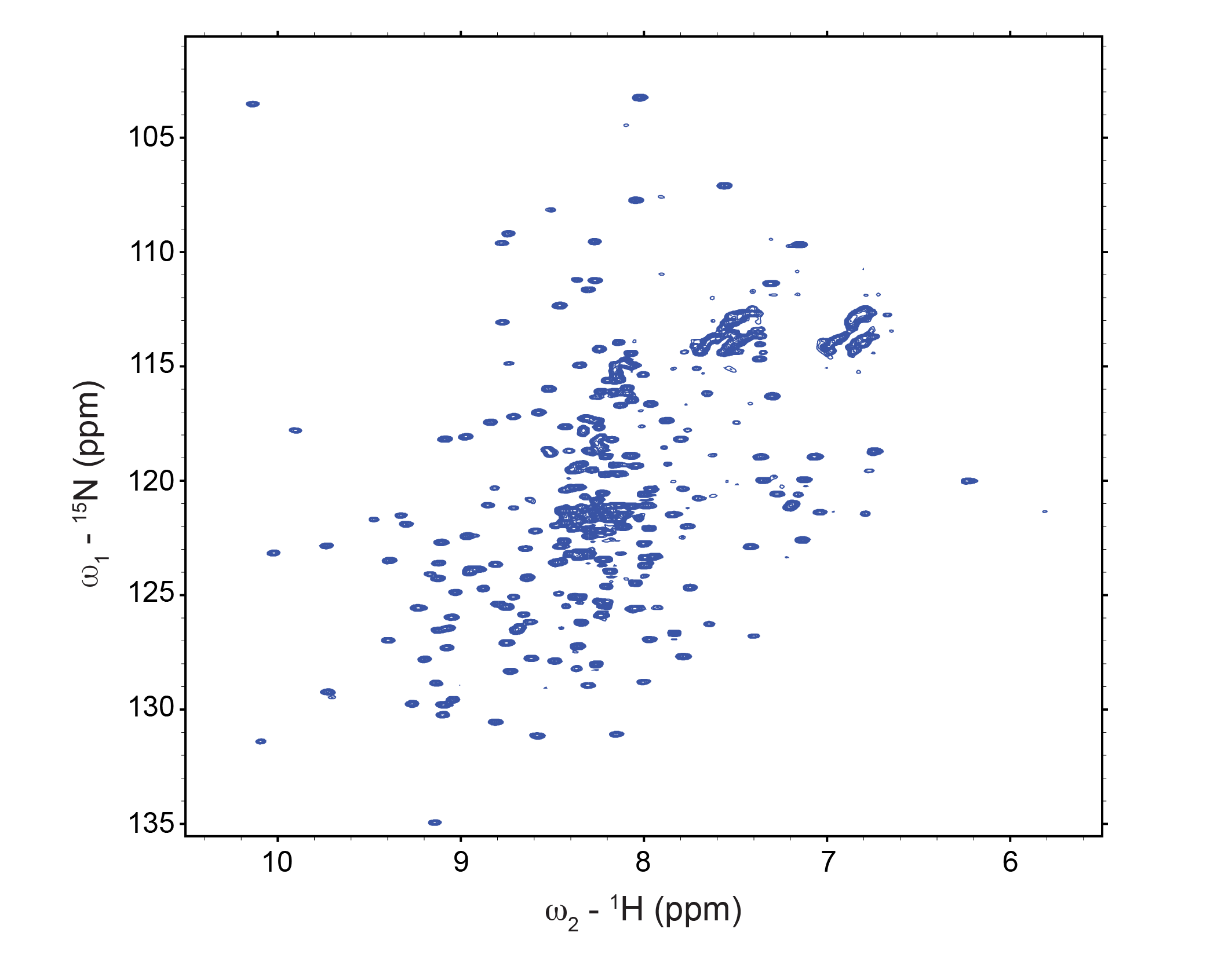

**Figure S2: ^15^N-^1^H TROSY-HSQC spectrum of apo-IsdH.**

The two-dimensional backbone HSQC spectrum of apo-IsdH^FL^ shows excellent peak dispersion for a protein construct of its size (66 kDa), that can be attributed to relative mobility between N1 and the extraction unit. Additionally, resonances corresponding to previously assigned backbone chemical shifts of both IsdH’s N1 domain and its N2N3 tri-domain extraction unit are present, supporting that the protein is correctly folded (11, 12). Additional signals at ~8 ppm in ^1^H spectrum can be attributed to the ~100 amino acid sequence that connects the N1 domain to the N2N3 tri-domain extraction unit.

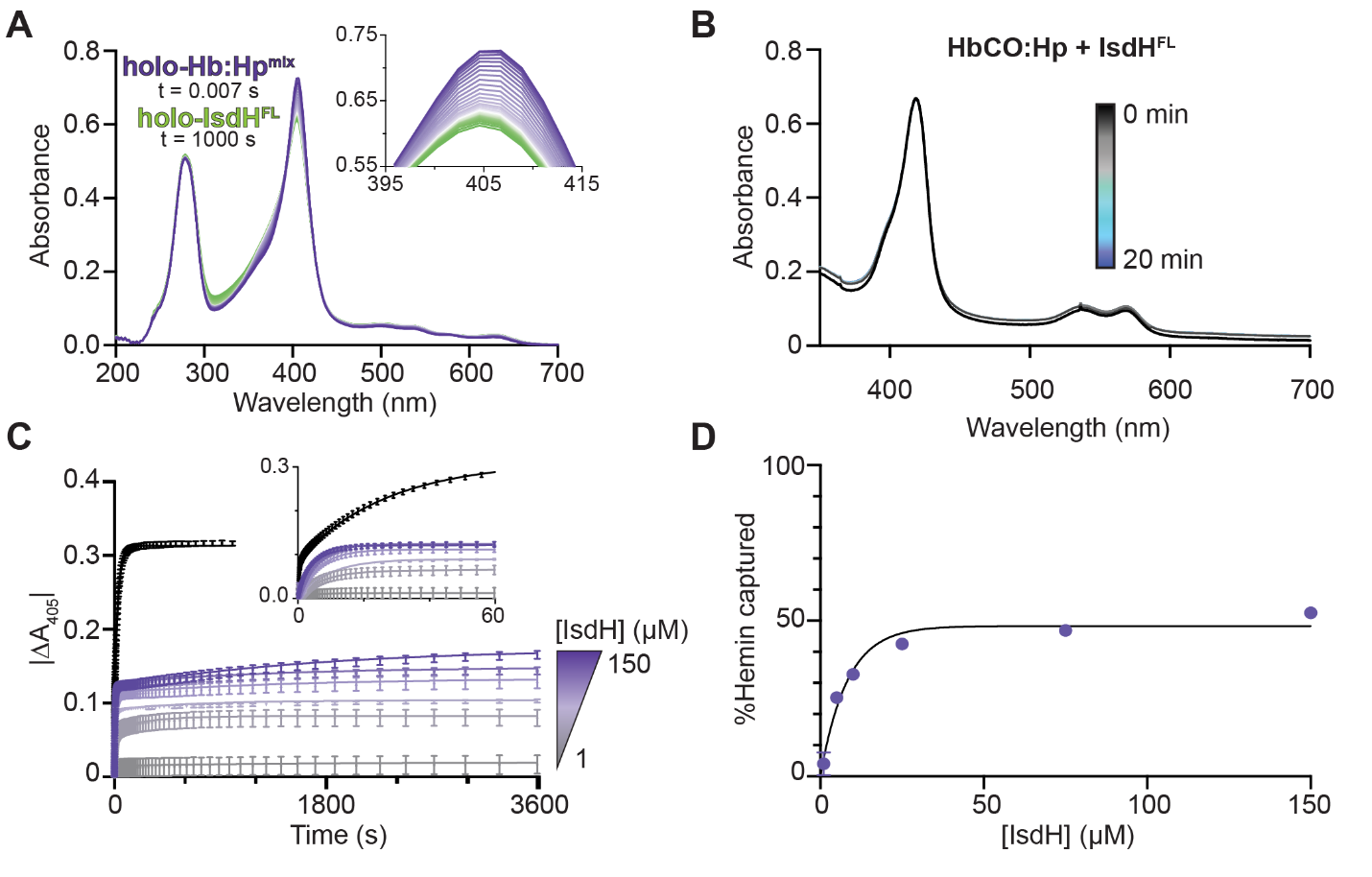

**Figure S3: Time resolved spectral changes of met- and carbonmonoxy-Hb:Hp (HbCO:Hp) in the presence of IsdH^FL^.**

(**A**) Time-dependent spectral changes that reflect hemin transfer from Hb:Hp^mix^ to IsdH^FL^. Traces are colored from t=0s (purple) to t=1000s (green). Inset: expanded spectral changes in the Soret region. (**B**) HbCO:Hp1-1 exhibits minimal spectral changes over a 20 minute timecourse in the presence of apo-IsdH^FL^. (**C**) Traces accompanying hemin transfer to IsdH^N2N3^ from Hb (black) and to IsdH^FL^ from Hb:Hp^mix^ (purple and gray). Inset: first 60s of hemin capture timecourse. (**D**) Percent of hemin captured from Hb:Hp^mix^ at 1-150 µM IsdH^FL^. Percent hemin captured was calculated as the ratio of |∆A_405_Hb:Hp|/|∆A_405_Hb|. Error bars corresponding to values determined by propagation of uncertainty are present, but they are too small to be visible for most data points at this scale.

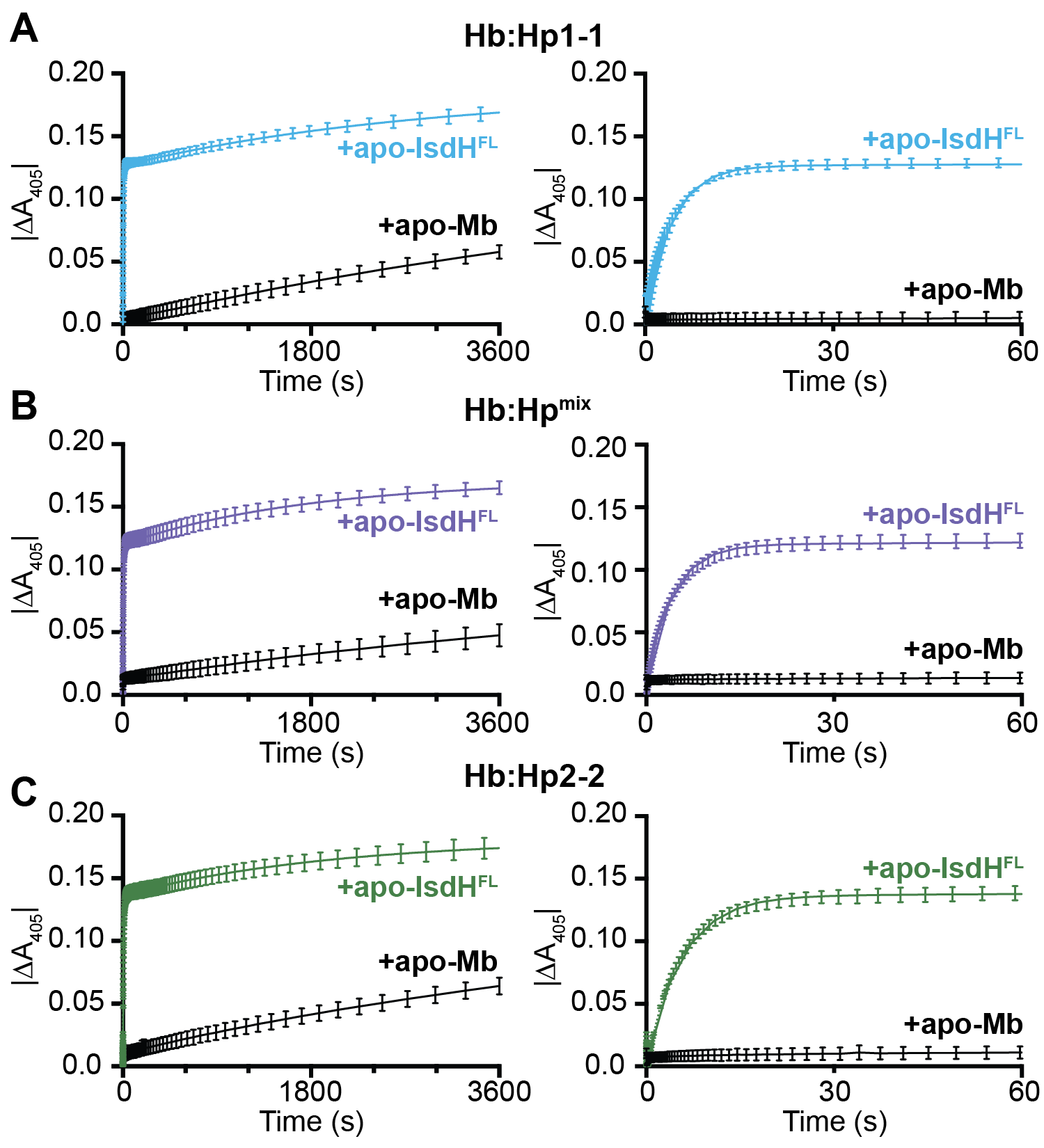

**Figure S4: Passive hemin release to apo-Mb and active hemin capture by apo-IsdH^FL^ across Hb:Hp phenotypes.**

Soret band absorbance changes accompanying passive hemin egress from metHb:Hp to 150 µM (30-fold excess) apo-Mb (black) and active hemin capture by 150 µM IsdH (**A**: blue, **B**: purple, and **C**: green) were monitored for 1 hour (left panels). Absorbance changes occurring in the first 60s of the reactions are shown in the right panels. Error bars represent the standard deviation of three replicates.

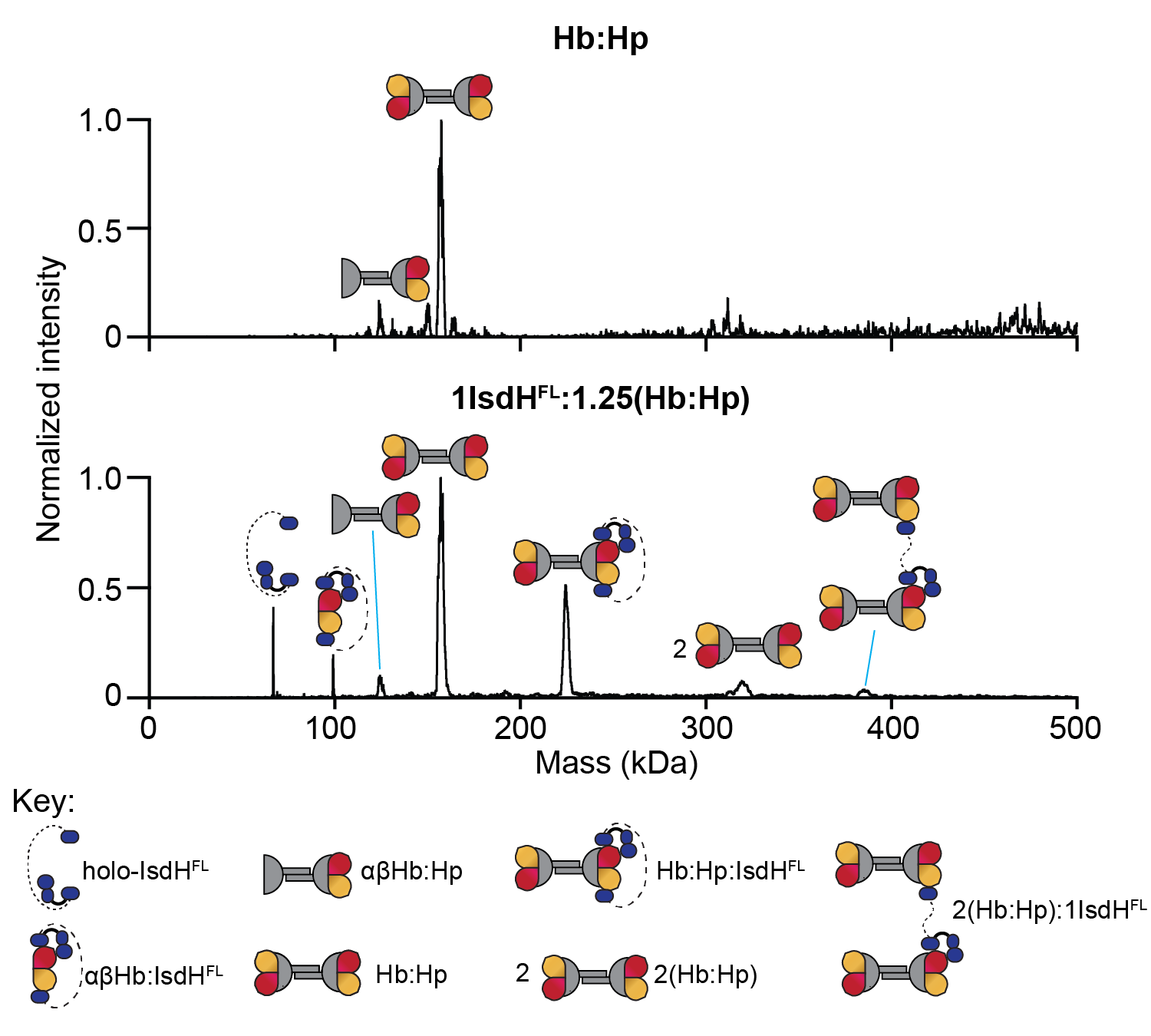

**Figure S5: Native mass spectrometry of the IsdH^FL^:Hb:Hp complex.**

Native mass spectra of (top) 1.25 µM Hb:Hp1-1 and (bottom) 1 µM apo-IsdH^FL^ + 1.25 µM Hb:Hp1-1 were acquired using electron capture charge reduction (ECCR). Hb:Hp-containing peaks are broadened due to microheterogeneity at the Hp glycosylation sites (13, 14). When Hb:Hp is combined with sub-stoichiometric apo-IsdH^FL^, we detect lone holo-IsdH^FL^, lone metHb:Hp, and complexes containing 1:1 IsdH^FL^:(Hb:Hp) and 1:2 IsdH^FL^:(Hb:Hp). The mass spectrum additionally contains species consistent with Hb:Hp fragmentation (IsdH^FL^ complexed to an Hb dimer and an Hp1-1 dimer bound to only one Hb dimer) and an Hb:Hp complex dimer. These species are likely either gas phase artifacts or are exceedingly scarce in solution, as they are not detected by SEC-MALS.

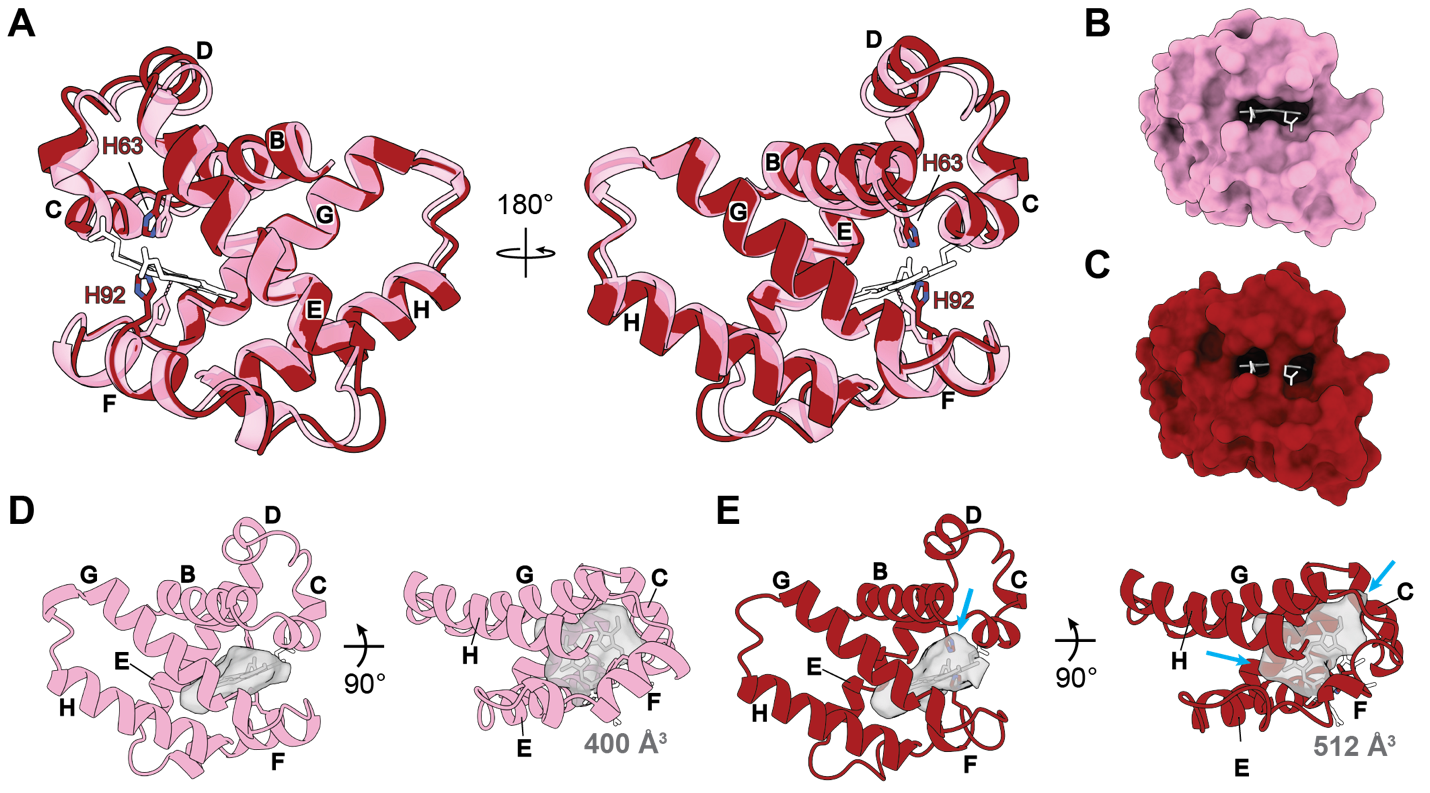

**Figure S6: Loss of βHb hemin within Hb:Hp is accompanied by substantial remodeling of the hemin pocket.**

(**A**) An overlay of βHb from the IsdH^FL^:Hb:Hp structure (red) and βHb from lone metHb (PDB: 6NBC, transparent pink) (15) reveals receptor-induced structural distortions to the hemin binding pocket. The distal and proximal histidines from each structure are shown as sticks and colored in correspondence with their parent chain. Hemin from the lone metHb structure is shown as white sticks with a black outline. The E- and F-helices, which contain the distal and proximal histidines (H63 and H92), shift inward toward the vacated hemin binding pocket, whereas the B-, C-, D-, and H- helices are all outwardly displaced. The A- and G-helix positions are largely unchanged. Residues N-terminal to the A-helix, within the A helix, and C-terminal to the H-helix are omitted for clarity. These structural rearrangements cause partial closure of the βHb hemin pocket to the exterior, which is evident upon comparing a surface representations of βHb within (**B**) lone metHb and (**C**) the IsdH^FL^:Hb:Hp complex. The hemin molecule from lone metHb is shown as white sticks in both panel (B) and panel (C). (**D**) Cartoon representation of lone metHb with its hemin shown as white sticks and its hemin binding pocket cavity shown as a gray surface. Residues N-terminal to the A-helix, within the A helix, and C-terminal to the H-helix are omitted for clarity. (**E**) βHb from the IsdH^FL^:Hb:Hp structure rendered as in (D). Hemin from an overlay with the lone metHb structure is shown as sticks to aid in visualizing the heme binding pocket. Areas where the hemin binding pocket has noticeably expanded are indicated with blue arrows in panel (E).

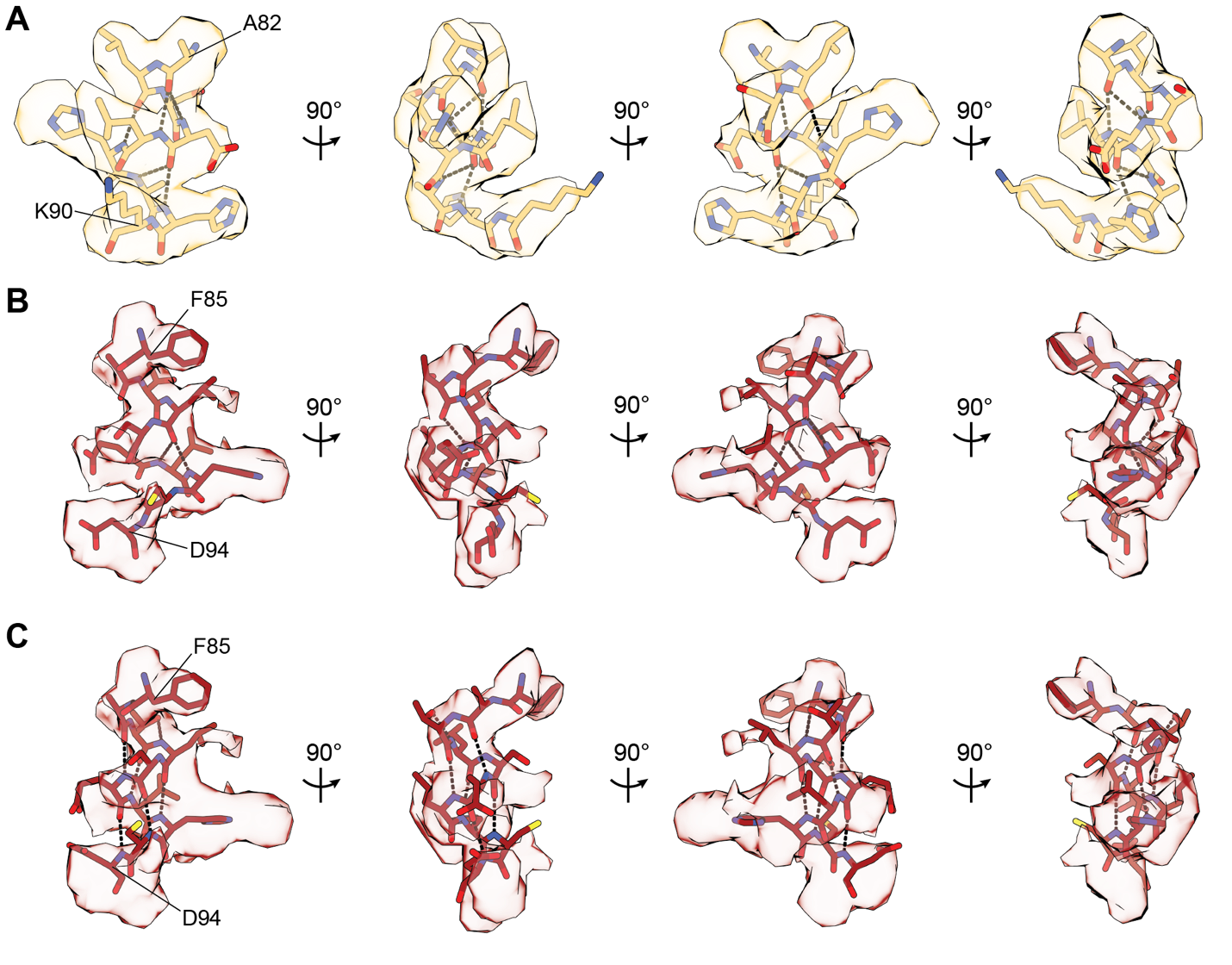

**Figure S7: F-helices of the IsdH^FL^:Hb:Hp complex.**

Cryo-EM map quality enabled placement of residues in the (A) αHb and (B) βHb F-helix. A loss of helicity is apparent in the βHb F-helix. During model building, we initially imposed helical restraints. However, the model to map fit for the βHb F-helix degrades when these restraints are enforced (C). Thus, the residues spanning the βHb F-helix in our model no longer satisfy the hydrogen bonding criteria for an α-helix. Hydrogen bonds are shown as black dashes.

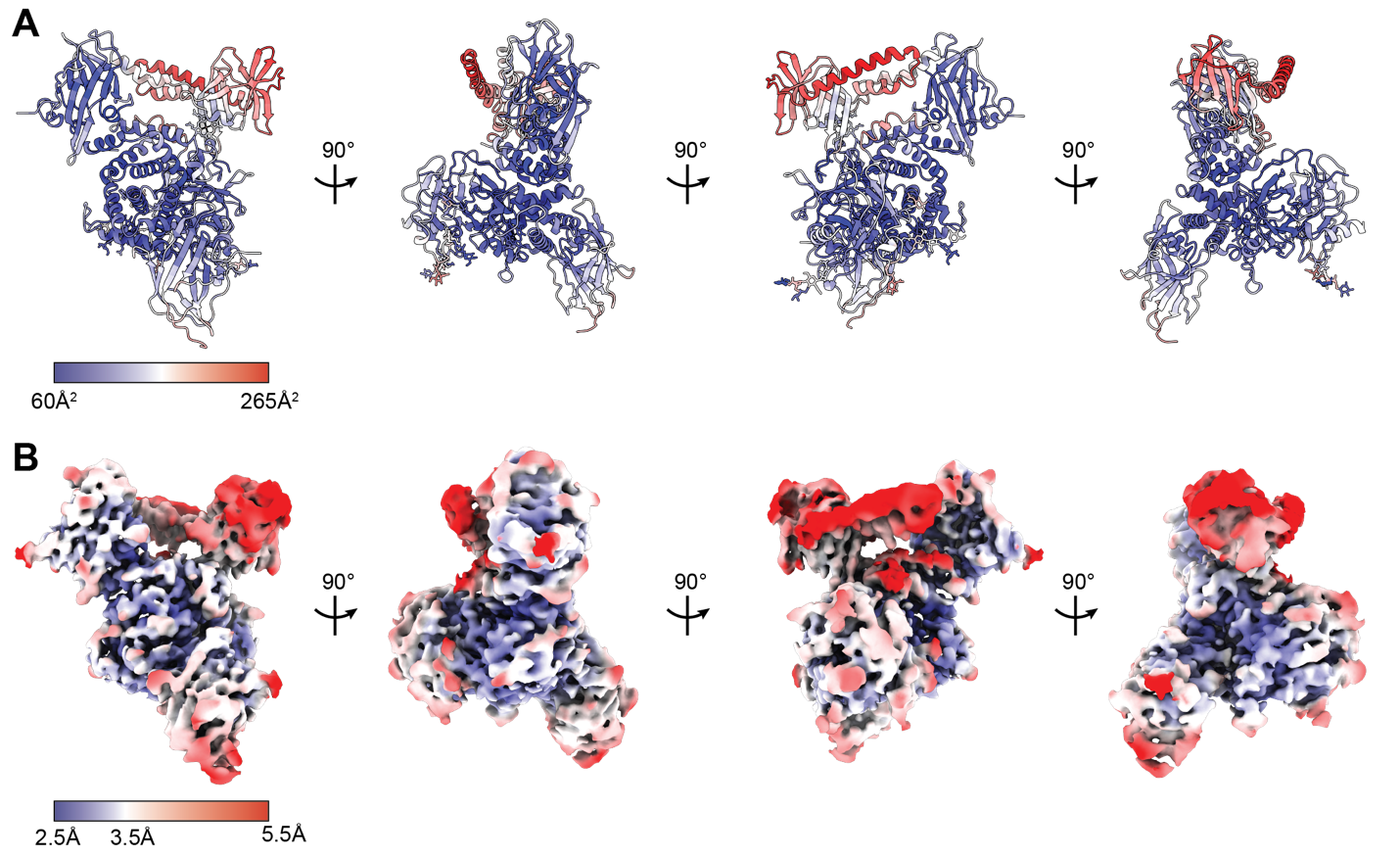

**Figure S8: B-factors and local resolution suggest conformational heterogeneity within the IsdH^FL^:Hb:Hp complex.**

(**A**) The IsdH^FL^:Hb:Hp complex exhibits B-factors ranging from ~60-265 Å^2^. Higher B-factors are observed in the IsdH L and N3 domains as well as the βHb F-helix. (**B**) These regions also have poor local resolution within the cryo-EM map.

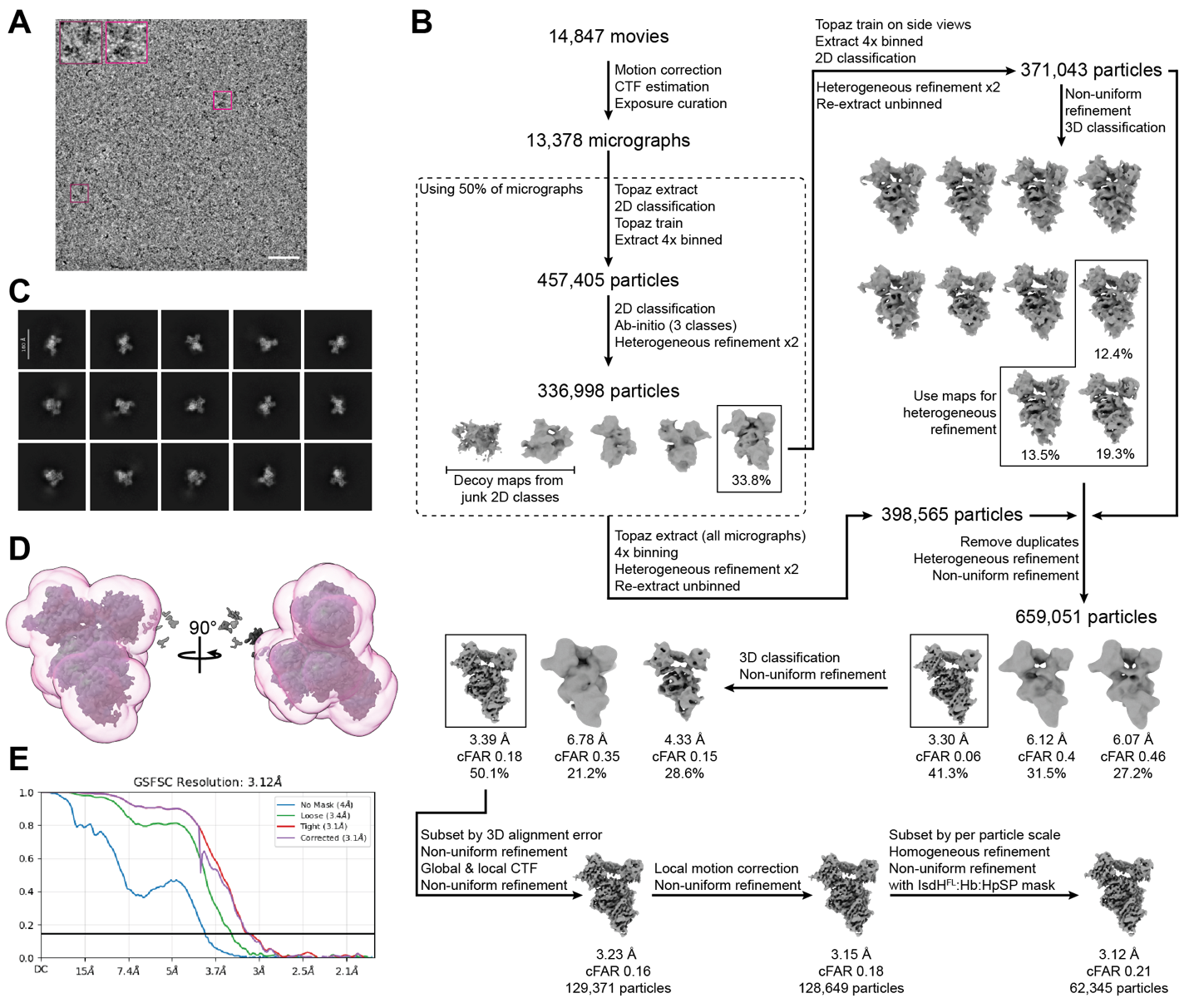

**Figure S9: Cryo-EM data processing.**

(**A**) Representative micrograph with IsdH^FL^:Hb:Hp particles boxed in pink and expanded in the inset. (**B**) Cryo-EM data processing workflow. (**C**) Representative 2D classes of the final particle stack. (**D**) Map from the penultimate refinement job (dark gray) in the mask used for the final non-uniform refinement (pink). (**E**) Gold-standard Fourier shell correlation curve with the 0.143 threshold shown as a horizontal black line.

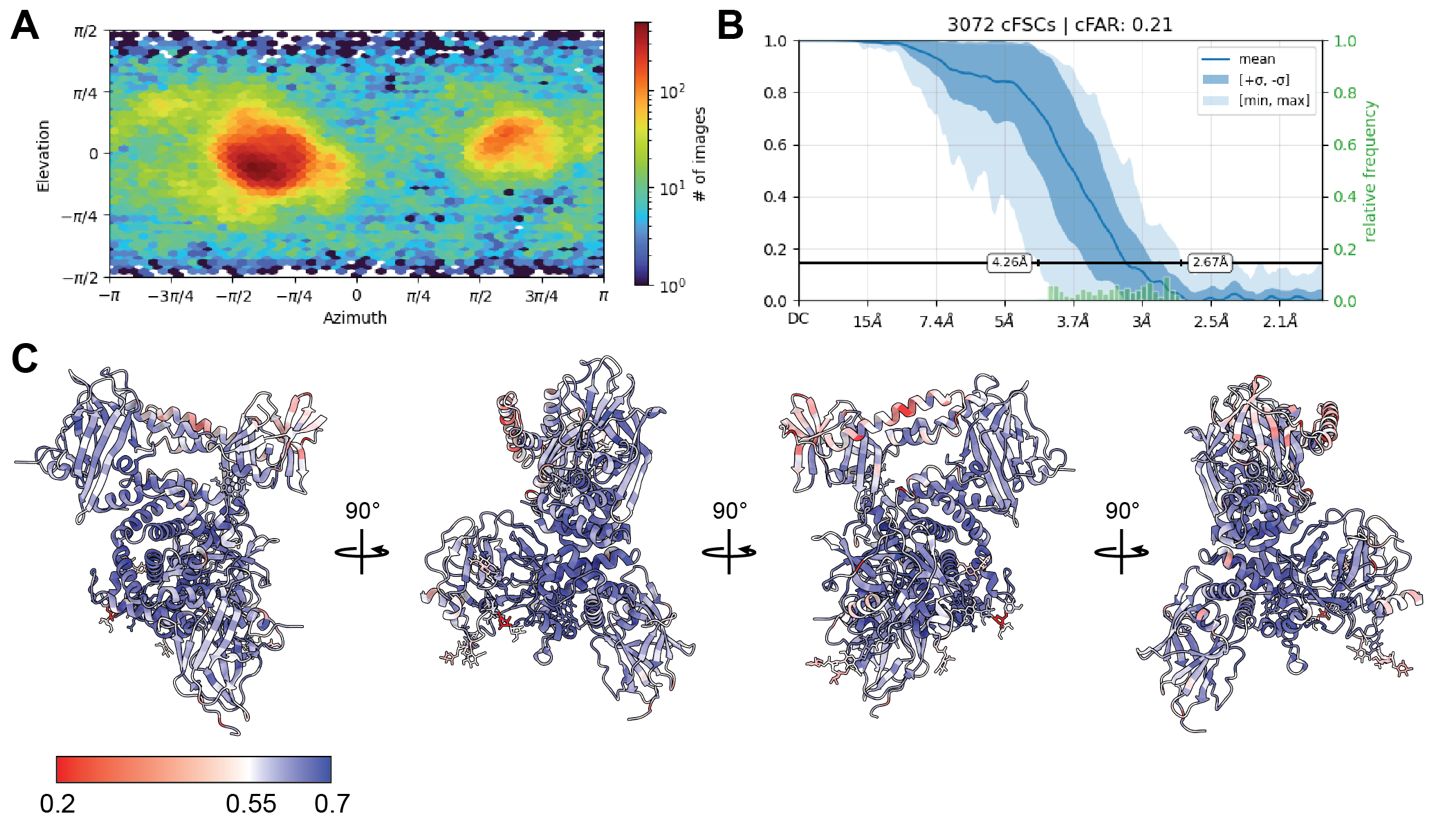

**Figure S10: Cryo-EM orientation diagnostics and model to map fit.**

The (**A**) particle viewing direction distribution and (**B**) conical FSCs (half-angle 20°) plots are indicative of preferred orientation. (**C**) The IsdH^FL^:Hb:Hp model colored by Q-score.

**Table S1. Kinetics of IsdH-mediated hemin extraction from hemoglobin:haptoglobin complexes.**

| **[IsdH] (µM)** | **Hemin source** | **ΔA_405_ (x 10^-1^)** | **% Hemin transferred** | **%Fast** | ***k*_fast_ (s^-1^) (x 10^-1^)** | ***k*_slow_ (s^-1^) (x 10^-3^)** | **Initial rate (µM s^-1^)** |
| --- | --- | --- | --- | --- | --- | --- | --- |
| 1 | Hb:Hp^mix^ | 0.127 ± 0.005 | 4% ± 3% | *n.d.** | *n.d.** | *n.d.** | *n.d.** |
| 5 |  | 0.797 ± 0.002 | 25.3% ± 0.3% | 78.0% ± 0.2% | 1.42 ± 0.01 | 3.3 ± 0.1 | 0.01 ± 0.04 |
| 10 |  | 1.0330 ± 0.0007 | 32.80% ± 0.09% | 87.95% ± 0.06% | 1.118 ± 0.002 | 1.37 ± 0.02 | 0.017 ± 0.001 |
| 25 |  | 1.340 ± 0.005 | 42.6% ± 0.4% | 85.2% ± 0.3% | 1.662 ± 0.005 | 0.63 ± 0.03 | 0.040 ± 0.005 |
| 75 |  | 1.476 ± 0.002 | 46.9% ± 0.2% | 85.1% ± 0.1% | 1.823 ± 0.003 | 0.74 ± 0.01 | 0.054 ± 0.002 |
| 150 |  | 1.655 ± 0.004 | 52.6% ± 0.2% | 68.9% ± 0.1% | 2.457 ± 0.005 | 0.561 ± 0.007 | 0.074 ± 0.003 |
| 150 | Hb:Hp1-1 | 1.82 ± 0.01 | 57.8% ± 0.6% | 67.5% ± 0.4% | 2.412 ± 0.007 | 0.350 ± 0.009 | 0.086 ± 0.008 |
| 150 | Hb:Hp2-2 | 1.788 ± 0.007 | 56.8% ± 0.4% | 74.7% ± 0.3% | 1.743 ± 0.003 | 0.50 ± 0.01 | 0.066 ± 0.005 |
| 150 (IsdH^N2N3^) | Hb | 3.15 ± 0.02 | 100.00% ± 0.09% | 28.40% ± 0.05% | 29.0 ± 0.2 | 36.71 ± 0.06 | 1.338 ± 0.001 |

**n.d.: At low IsdH concentrations hemin transfer was negligible and could not be fit to an equation.*

Kinetic parameters characterizing hemin capture from Hb or Hb:Hp in the presence of apo-IsdH^FL^ or apo-IsdH^N2N3^. Values were determined by monitoring the reduction in A405 after mixing of 5 µM Hb or Hb:Hp on a hemin basis with 1, 5, 10, 25, 75, or 150 µM IsdH. Reactions were fit to a biphasic exponential equation. The percentage of hemin transfer (% Hemin transferred) was calculated as the ratio of |∆A_405_Hb:Hp|/|∆A_405_Hb|. %Fast is a parameter determined by curve fitting that describes the portion of heme loss attributed to the fast phase of the reaction (i.e., loss from βHb). The initial rate of heme release in µM s^-1^ was calculated by multiplying the starting hemin concentration (5 µM) by the first derivative of the exponential decay equation to which each dataset was fit.

**Table S2: Buried surface area analysis of βHb, the IsdH extraction unit, and βHb-derived hemin across structures capture at various stages of hemin extraction.**

|  | **IsdHN2N3:βHb** | **IsdHN3:βHb** | **βHb:heme/hemin** | **IsdHN3:heme/hemin** |
| --- | --- | --- | --- | --- |
| pre-extraction (PDB: 9S3P) | 1235.6 Å^2^ | 507.7 Å^2^ | 546.9 Å^2^ | 77.7 Å^2^ |
| mid-extraction (PDB: 6TB2) | 1103.9 Å^2^ | 424.6 Å^2^ | 412.3 Å^2^ | 276.7 Å^2^ |
| post-extraction (this work) | 860.1 Å^2^ | 207.6 Å^2^ | 201.9 Å^2^ | 441.8 Å^2^ |

Analysis was performed using the PISA server (16) on the IsdHN2N3, βHb, and heme/hemin interfaces across the post-extraction complex (this work) and structures of the isolated extraction unit bound to HbCO (PDB: 9S3P) or to Hb:HpSP (an Hb dimer bound to recombinant, non-glycosylated Hp SP domain, PDB: 6TB2) (17, 18). The IsdHN2N3:βHb interface buried surface area decreases as hemin transfer progresses, with most of the reduction attributable to movement of the N3 domain. As hemin moves to IsdHN3, its buried surface area decreases with βHb and increases with IsdHN3.

**Table S3: Cryo-EM data collection, image analysis, modelling, refinement, and validation statistics.**

| **Dataset** | **IsdH^FL^ + metHb:Hp1-1 (PDB: 36EB, EMD-77412)** |
| --- | --- |
| **Map refinement** | Focused on IsdH N1, IsdH N2N3, αβHb, and HpSP |
| **Data collection and processing** | |
| **Microscope** | UCLA Titan Krios G4 |
| **Voltage (kEV)** | 300 |
| **Detector** | Falcon 4i |
| **Nominal magnification** | 130,000x |
| **Data acquisition software** | EPU |
| **Electron dose (e^-^/Å^2^)** | 50 |
| **Pixel size (Å)** | 0.97 |
| **Defocus range (µM)** | -1.0 to -2.5 |
| **Number of movies collected** | 14,847 |
| **Number of micrographs used** | 13,378 |
| **Reconstruction** | |
| **3D processing package** | CryoSparc |
| **Initial particle images (no.)** | 62,345 |
| **Final particle images (no.)** | 457,405 |
| **Symmetry** | C1 |
| **FSC (0.143)** | 3.12 |
| **Refinement** | |
| **Initial model** | Alphafold 3 |
| **Model refinement package** | Phenix |
| **Non-hydrogen atoms** | 8,372 |
| **Protein residues** | 1024 |
| **R.M.S. deviations** |  |
| **Bond lengths (Å)** | 0.002 |
| **Bond angles (°)** | 0.506 |
| **Validation** | |
| **MolProbity score** | 1.49 |
| **Clashscore** | 3.75 |
| **Poor rotamers (%)** | 1.46 |
| **Ramachandran (%)** |  |
| **Favored** | 96.25 |
| **Allowed** | 3.75 |
| **Disallowed** | 0 |
| **Fit to map (CC_mask_)** | 0.81 |
| **Q-score** | 0.462 |

**Supplementary references**

1.	T. Bepler, *et al.*, Positive-unlabeled convolutional neural networks for particle picking in cryo-electron micrographs. *Nat. Methods* **16**, 1153–1160 (2019).

2.	A. Punjani, D. J. Fleet, 3D variability analysis: Resolving continuous flexibility and discrete heterogeneity from single particle cryo-EM. *J. Struct. Biol.* **213**, 107702 (2021).

3.	M. D. Tyka, *et al.*, Alternate States of Proteins Revealed by Detailed Energy Landscape Mapping. *J. Mol. Biol.* **405**, 607–618 (2011).

4.	K. I. P. Le Huray, *et al.*, To 200,000 *m* / *z* and Beyond: Native Electron Capture Charge Reduction Mass Spectrometry Deconvolves Heterogeneous Signals in Large Biopharmaceutical Analytes. *ACS Cent. Sci.* **10**, 1548–1561 (2024).

5.	M. T. Marty, *et al.*, Bayesian Deconvolution of Mass and Ion Mobility Spectra: From Binary Interactions to Polydisperse Ensembles. *Anal. Chem.* **87**, 4370–4376 (2015).

6.	J. Marley, M. Lu, C. Bracken, A method for efficient isotopic labeling of recombinant proteins. *J. Biomol. NMR* **20**, 71–75 (2001).

7.	K. Pervushin, R. Riek, G. Wider, K. Wuthrich, Attenuated T2 relaxation by mutual cancellation of dipole–dipole coupling and chemical shift anisotropy indicates an avenue to NMR structures of very large biological macromolecules in solution. *Proc. Natl. Acad. Sci. U.S.A.* **94**, 12366–12371 (1997).

8.	C. B. F. Andersen, *et al.*, Haptoglobin. *Antioxid. Redox Signal.* **26**, 814–831 (2017).

9.	A. di Masi, *et al.*, Haptoglobin: From hemoglobin scavenging to human health. *Mol. Asp. Med.* **73**, 100851 (2020).

10.	M. S. Hargrove, *et al.*, His64(E7)–>Tyr apomyoglobin as a reagent for measuring rates of hemin dissociation. *J. Biol. Chem.* **269**, 4207–4214 (1994).

11.	R. M. Pilpa, R. T. Clubb, NMR Resonance Assignments of the NEAT (NEAr Transporter) Domain from the Staphylococcus aureus IsdH Protein. *J. Biomol. NMR* **33**, 137 (2005).

12.	T. Spirig, R. T. Clubb, Backbone 1H, 13C and 15N resonance assignments of the 39 kDa staphylococcal hemoglobin receptor IsdH. *Biomol. NMR Assign.* **6**, 169–172 (2012).

13.	D. Wu, W. B. Struwe, D. J. Harvey, M. A. J. Ferguson, C. V. Robinson, N-glycan microheterogeneity regulates interactions of plasma proteins. *Proc. Natl. Acad. Sci. U.S.A.* **115**, 8763–8768 (2018).

14.	S. Tamara, V. Franc, A. J. R. Heck, A wealth of genotype-specific proteoforms fine-tunes hemoglobin scavenging by haptoglobin. *Proc. Natl. Acad. Sci. U.S.A.* **117**, 15554–15564 (2020).

15.	M. A. Herzik, M. Wu, G. C. Lander, High-resolution structure determination of sub-100 kDa complexes using conventional cryo-EM. *Nat. Commun.* **10**, 1032 (2019).

16.	E. Krissinel, K. Henrick, Inference of Macromolecular Assemblies from Crystalline State. *J. Mol. Biol.* **372**, 774–797 (2007).

17.	V. B. Comani, *et al.*, Hemoglobin receptor redundancy in Staphylococcus aureus: molecular flexibility as a determinant of divergent hemophore activity. *J. Struct. Biol. X* **12**, 100138 (2025).

18.	J. H. Mikkelsen, K. Runager, C. B. F. Andersen, The human protein haptoglobin inhibits IsdH-mediated heme-sequestering by Staphylococcus aureus. *J. Biol. Chem.* **295**, 1781–1791 (2020).
